# Prostaglandin D2 pathway in a transgenic rat model of Alzheimer’s disease: therapeutic potential of timapiprant a DP2 antagonist

**DOI:** 10.1101/2022.03.30.486444

**Authors:** Charles H. Wallace, Giovanni Oliveros, Peter A Serrano, Patricia Rockwell, Lei Xie, Maria Figueiredo-Pereira

**Author notes:** **Corresponding Author:** Maria E. Figueiredo-Pereira, Department of Biological Sciences, Hunter College and Graduate Center, City University of New York, 695 Park Ave., New York, NY 10065, USA.

## Abstract

The cyclooxygenase pathway, a key mediator of inflammation, is implicated in Alzheimer’s disease (AD). A deeper investigation is required into the contributions of this pathway to the neuropathology of AD. Cyclooxygenases produce prostaglandins, which have multiple receptors and functions including inflammation, nociception, sleep, cardiovascular maintenance and reproduction. In the brain, prostaglandin D2 (PGD2) is the most abundant prostaglandin, increases the most under pathological conditions, and plays roles in sleep, stroke and inflammation. PGD2 signals through its DP1 and DP2 receptors and their activation can be protective or detrimental. We address the relationship between the PGD2 pathway and AD neuropathology with F344-AD transgenic (Tg-AD) rats that exhibit age-dependent and progressive pathology similar to AD patients. We analyzed the PGD2 pathway in the hippocampus of wild type (WT) rats and their Tg-AD littermates, at the age of 11 months, when Tg-AD rats exhibit plaques and perform significantly worse in hippocampal-dependent cognitive tasks than WT rats. Using mass spectrometry, we determined that PGD2 levels were at least 14.5-fold higher than PGE2, independently of genotype. Immunohistochemistry established that microglial DP1 receptors were more abundant and neuronal DP2 receptors were fewer in Tg-AD than in WT rats. RNA sequencing profiling of 33 genes involved in the PGD2 and PGE2 pathways revealed that mRNA levels were the highest for L-PGDS, the major PGD2 synthase in the brain. To evaluate the pathophysiological significance of our findings on the PGD2 pathway, we treated a subset of rats (WT and Tg-AD males) with timapiprant, a potent and highly selective oral DP2 antagonist being developed as a once-daily oral treatment in patients with allergic inflammation. We conclusively show that timapiprant significantly mitigated some of the AD pathology exhibited by the Tg-AD male rats. More comprehensive studies are necessary to support the therapeutic potential of timapiprant and that of other PGD2-related compounds in the treatment of AD.

## 1. Introduction

Alzheimer’s disease (AD) is the most common type of dementia, is highly prevalent in the ageing population, and will become more prevalent as life expectancy continues to rise. AD is a multifactorial disease, and chronic neuroinflammation is recognized as a critical factor in its pathogenesis [12]. A major player in inflammation is the cyclooxygenase (COX) pathway [36]. The COX pathway generates prostaglandins (PGs), which are bioactive signaling lipids responsible for many processes including inflammation [8]. PG signaling is implicated in AD, as some PGs aggravate its pathology while others may remediate it (reviewed in [10]). Based on data from epidemiological studies, there is a decreased risk of AD in patients taking NSAIDs, which are inhibitors of the COX pathway [66]. Inhibiting COXs with NSAIDs was proposed to be a promising therapeutic strategy. However, while long-term use of NSAIDs is associated with a reduced incidence of AD in epidemiologic studies, randomized controlled trials did not replicate these findings [17]. Moreover, NSAIDs target COX-1 and/or COX-2 enzymes stopping most PG synthesis. Non-specific inhibition of PG synthesis can have a variety of negative side effects, because PGs have many functions including inflammation, nociception, sleep, cardiovascular maintenance and reproduction [44]. Accordingly, negative side effects such as renal failure, heart problems, and stroke were reported for NSAIDs during these clinical trials [17]. Thus, NSAIDs cannot currently be recommended for either primary prevention or treatment of AD. Based on these concerns, it is important to find new targets further downstream in the COX pathway, such as specific PG signaling that can be explored for potential therapeutic intervention.

In the current study, we investigated the relationship between the prostaglandin D2 (PGD2) pathway and AD neuropathology. PGD2 is the most abundant prostaglandin in the brain, and is the one that increases most under neuropathological conditions [1, 35]. In the brain, lipocalin-type prostaglandin D synthase (L-PGDS) is the primary synthase for PGD2 [64]. PGD2 undergoes a nonenzymatic dehydration producing PGJ2 (reviewed in [19]). PGD2 signals through its two antagonistic receptors, DP1 and DP2, the latter also known as CRTH2 or GPR44. DP1 activation by PGD2 is coupled to the G protein G_s_ leading to an increase in cAMP and PKA activation (reviewed in [68]). DP1 activation also elicits an increase in calcium, but the latter is cAMP-dependent (reviewed in [68]). DP1 plays a well characterized role in sleep function [5], and in vivo studies show that DP1 modulation is protective in ischemic and hemorrhagic models of stroke [3–5, 18]. DP2 receptor activation by PGD2 and PGJ2, is coupled to the G protein G_i_ leading to a decrease in cAMP and an increase in calcium mobilization, both of which can lead to neuronal damage (reviewed in [68]). For example, *in vitro* studies show adverse outcomes when treating hippocampal neuronal cultures and organotypic slices with DP2 agonists [34, 35].

The PGD2 pathway is thoroughly studied in diseases with airway inflammation and reproduction [40, 54], but its role in AD neuropathology remains unclear. Investigating the relationship between the PGD2 pathway and AD neuropathology is important as it could lead to new therapeutic strategies to treat neuroinflammation in pre or early stages of AD, and slow down its neuropathology. Our studies focus on the PGD2 pathway in conjunction with neuronal loss and microgliosis, both known to occur in AD. Microglia are the resident macrophages of the brain, and have three different morphologies based on their function and shape [31]. The majority of microglia are *ramified* with long slender processes, and play a role in surveillance. *Reactive* microglia, present in intermediate numbers, exhibit shorter processes and a larger soma than ramified microglia, and are in an activated state producing immune modulators. Finally, *amoeboid* microglia are the fewest, have the largest soma and fewest processes, and perform phagocytosis. The relation between PGD2 signaling and microglia morphology remains to be established.

We investigate the PGD2 pathway in the hippocampus of a transgenic rat model of AD, the TgF344-AD (Tg-AD) rats and their wild type (WT) littermates. Tg-AD rats express the Swedish mutation (KM670/671NL) of human amyloid precursor protein (APPswe), and the Δ exon 9 mutation of human presenilin-1 (PS1ΔE9), both driven by the prion promoter [13]. Tg-AD rats develop AD neuropathology including cerebral amyloidosis, tauopathy, gliosis, and neuronal loss, as well as cognitive deficits, all in a progressive age-dependent manner.

To explore whether targeting PGD2 pathway has therapeutic potential for AD, we chose to treat a subset of rats (WT and Tg-AD males) with timapiprant (also known as OC000459), a potent and highly selective oral DP2 antagonist. Timapiprant is an indole-acetic acid derivative that potently displaces [3H]PGD2 from human recombinant DP2 (Ki = 0.013 μM), rat recombinant DP2 (Ki = 0.003 μM), and human native DP2 (Th2 cell membranes (Ki = 0.004 μM) [51]. Moreover, timapiprant does not interfere with the ligand binding properties or functional activities of other prostanoid receptors (EP1-4 receptors, DP1, thromboxane receptor, prostacyclin receptor, and prostaglandin F receptor) [51]. Timapiprant, which seems to be safe and well tolerated, is being developed for oral treatment of patients with allergic inflammation in diseases such as asthma and allergic rhinitis [41]. Many DP2 antagonists attenuate the inflammatory response in animal studies for these diseases, some demonstrate efficacy in phase II studies in adults with asthma, and several phase III trials are evaluating the long-term safety and efficacy of these drugs in adult and pediatric patients with moderate-to-severe asthma [41].

In summary, our analyses compared PGD2, PGE2, PGJ2 and thromboxane B2 concentrations, the cellular distribution of the DP1 and DP2 receptors, mRNA profiles for 33 genes involved in the PGD2 and PGE2 pathways, Aβ plaque burden, neuronal loss, and microgliosis in the hippocampus of 11 month old WT versus Tg-AD rats, as well as their cognitive performance. Our findings are the first to comprehensively address various aspects of the PGD2 pathway in a rat model of AD. We establish that PGD2 levels in the hippocampus are at least 14.5-fold higher than those for PGE2, independently of genotype. In addition, our data reveal significant differences in DP1 and DP2 receptor levels, respectively in microglia and neurons of Tg-AD rats compared to controls. Our transcriptome assessment identified L-PGDS as the most abundant mRNA of the 33 genes analyzed. Notably, we established that the DP2 antagonist timapiprant ameliorated the AD pathology developed by Tg-AD male rats. Overall, our studies provide novel insights for the development of therapeutics that target the PGD2 pathway to treat neuroinflammation in AD.

## 2. Materials and Methods

### 2.1. TgF344-AD transgenic rat model of AD

Fisher transgenic F344-AD (Tg-AD) rats [13] express human Swedish amyloid precursor protein (APPswe) and Δ exon 9 presenelin-1 (PS1ΔE9) driven by the prion promoter, at 2.6- and 6.2-fold higher levels respectively, than the endogenous rat proteins [13]. We purchased the Tg-AD rats and their WT littermates from Rat Resource and Research Center (RRRC, Columbia, MO) at four weeks of age. The rats were housed in pairs upon arrival and maintained on a 12h light/dark cycle with food and water available *ad libitum*. The Institutional Animal Care and Use Committee at Hunter College approved all animal procedures.

The Tg-AD rats exhibit a progressive age-dependent AD-like pathology as depicted in Fig. 1a and described in [13], including cognitive deficits, neuronal loss, Aβ plaque and neurofibrillary tangle burden, as well as gliosis. No differences in pathology was reported between sexes [13].

**Fig. 1.**
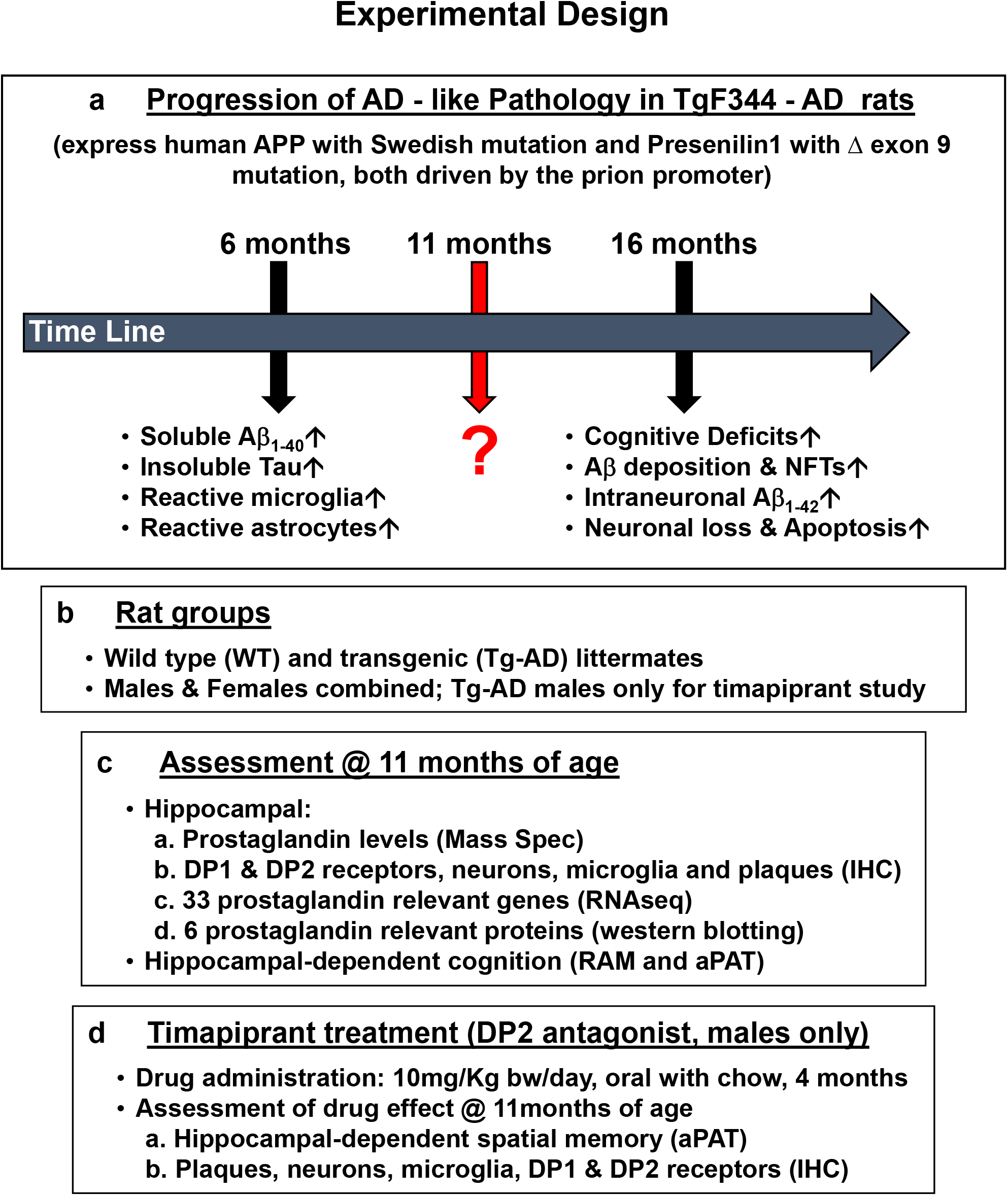
Schematic representation of the experimental design **a** Time line of the progression of the AD pathology developed by Tg-AD rats. **b** Rat groups used in the study. **c** Assessments of the AD pathology developed by the Tg-AD rats at 11 months of age. **d** Timapiprant treatment overview.

### 2.2. Experimental Design

A total of 93 rats for the combined female and male studies [WT n = 49 (27 females, 22 males), Tg-AD n = 44 (25 females, 19 males)] across multiple cohorts were used (Fig. 1b). For the timapiprant-treated studies, 9 WT and 9 Tg-AD males were used. At seven months, Tg-AD and WT rats began timapiprant treatment (cat # HY-15342, MCE, Monmouth Junction, NJ) with 10 mg/kg body weight/day/rat administered orally in rodent chow (Research Diets Inc. NJ) for 4 months. Thus, rats were sacrificed at 11 months of age. Future studies will include timapiprant-treated females.

We evaluated all rats at 11 months of age as described in Fig. 1c and 1d. Hippocampal-dependent cognitive deficits were estimated with the 8-Arm radial arm maze (RAM), which is a passive behavioral task, and/or the active-place avoidance task (aPAT). Following behavioral testing, the rats were sacrificed, the brains rapidly isolated and bisected into hemispheres, and processed for the different assays as described below and in Fig. 1c and 1d.

### 2.3. Tissue collection and preparation

At 11-months of age, the rats were anesthetized with an intraperitoneal injection containing ketamine (100 mg/ kg body weight) and xylazine (10 mg/kg body weight), and then transcardially perfused with chilled RNAase free PBS. The brain left hemispheres were micro-dissected into different regions, snap frozen with a CoolRack over dry ice, and the hippocampi tissue used for mass spectrometry, RNAseq, or western blot analyses. Whole right brain hemispheres were placed in a 4% paraformaldehyde/PBS solution for 48 hours at 4°C, followed by cryoprotection with a 30% sucrose/PBS solution to prevent water-freeze damage, and then flash frozen using 2-methylbutane, and stored at −80°C until sectioning for IHC.

### 2.4. LC-MS/MS for PG quantification

Rat hippocampal tissue from 11-month WT (n = 31) and Tg-AD (n = 32) rats were analyzed by quantitative LC-MS/MS to determine PGD2, PGE2, PGJ2 and thromboxane B2 (TxB2) concentrations using the standard calibration curves of each compound. Samples were prepared as previously described [7]. In summary, hippocampal tissues were homogenized in PBS using a BeadBug microtube homogenizer, then a 10-mg wet weight equivalent of homogenate was removed and further diluted 1:1 with 1% formic acid. Deuterated internal standards were added and loaded on a Biotage SLE+ cartridge, and were eluted twice with t-butlymethylether. The eluent was spiked with a trap solution consisting of 10% glycerol in methanol with 0.01 mg/ml butylated hydroxytoluene. Samples were dried in a speed vacuum at 35°C, the tubes were washed with hexane and re-dried. The residue was dissolved in 80:20 water:acetonitrile with butylated hydroxytoluene and spin filtered with a 0.22 μm Millipore Ultrafree^®^ filter. 30 μl of sample were analyzed. Prostaglandin standard curves were spiked into PBS and prepared identically to the samples. Area ratios were plotted and unknowns determined using the slopes.

PGs were analyzed using a 5500 Q-TRAP hybrid/triple quadrupole linear ion trap mass spectrometer (Applied Biosystems, Carlsbad, CA) with electrospray ionization (ESI) in negative mode as previously described [7]. The mass spectrometer was interfaced to a Shimadzu (Columbia, MD) SIL-20AC XR auto-sampler followed by 2 LC-20AD XR LC pumps. The scheduled MRM transitions were monitored within a 1.5 min time-window. Optimal instrument parameters were determined by direct infusion of each analyte. The gradient mobile phase delivered at a flow rate of 0.5 ml/min, consisted of two solvents, 0.05% acetic acid in water and acetonitrile. The analytes were resolved on a Betabasic-C18 (100×2 mm, 3 μm) column at 40°C using the Shimadzu column oven. Data were acquired using Analyst 1.5.1 and analyzed using Multiquant 3.0.1(AB Sciex, Ontario, Canada).

### 2.5. Immunohistochemistry

Coronal sections were sliced into 30 μm sections using a cryostat (Lecia CM3050 S). IHC was restricted to dorsal hippocampal tissue within the following Bregma coordinates: −3.36 mm to −4.36mm [50]. Sections were mounted on gelatin slides and immunostained as previously described [6]. Following immunostaining, a mounting media of VectaShield® with DAPI (Vector Labs # H-1200-10) was used and slides were stored in the dark at 4°C until imaged. Sections were viewed on a Zeiss Axio Imager M2 with AxioVision software to capture ZVI files of 10x and 20x mosaic images of the whole hippocampus, and then converted to TIF files. Signal density (O.D.) was quantified using Image J as previously described [14].

Two to three sections (averaged) from each rat were immunostained with either a combination of anti-DP1 and anti-Iba1 antibodies or anti-DP2 and anti-NeuN antibodies. Primary and secondary antibodies are listed in Supplemental Table 10. For quantification the following thresholds were used: *DP1*: mean + 1.5*std, particles analyzed were in the range: 10-10000, and circularity: 0-1.00; *Iba1*. mean + 1.5*std, particles analyzed were in the range: 50-8000, and circularity: 0-1.00; *DP2*: mean + 1.5*std, particles analyzed were in the range: 10-5000, and circularity: 0-1.00; *NeuN*: mean + 1.5*std, particles analyzed were in the range: 10-10000, and circularity: 0-1.00.

Colocalization was analyzed by measuring the overlap of the masks for the two channels. For DP1 and Iba1 we report the number of microglia co-localized with DP1 per specific area (nm^2^). For DP2 and NeuN we report the percentage DP2 and NeuN signals co-localized within a specific area (nm^2^). Additionally, Iba1+ ramified, reactive, and amoeboid microglia phenotypes were analyzed for circularity based on the ImageJ form factor (FF = 4π x area/perimeter^2^): ramified (FF < 0.50), reactive (FF: 0.50 to 0.70), and amoeboid (FF>0.70) [14].

Astrocytes were immunostained with an anti-GFAP antibody as listed in Supplemental Table 10. For quantification, the following thresholds were used: mean + 1.5*std, particles analyzed were in the range: 30-1000, and circularity 0-1.00.

### 2.6. RNAseq analysis

Hippocampal tissue was used for RNAseq analysis outsourced to the UCLA Technology Center for Genomics & Bioinformatics services. Samples from five male WT and five male Tg-AD rats were compared, and the same for female rats. Briefly, total RNA was isolated from the hippocampal tissue using the RNeasy Mini Kit from Qiagen. The integrity of total RNA was examined by the Agilent 4200 TapeStation System. Libraries for RNAseq were constructed with the Kapa Stranded mRNA Kit (Roche, cat. KK8421) to generate strand-specific RNAseq libraries, which were amplified and sequencing was performed with the HiSeq3000 sequencer. Gene expression data were normalized as reads per million (RPM) using the TMM method. Differentially expressed genes between WT and Tg-AD rats for each sex were determined using the edgeR program [53]. RPMs were analyzed for fold-change, *p* values, and FDR for each gene (Supplemental Table 8).

### 2.7. Western blot analysis

Hippocampal tissue (20-25 mg) was homogenized in TBS for 90 sec at 25°C with the Bedbug Microtube Homogenizer (3,400 rpm, Model D1030, Benchmark Scientific). The supernatant was stored for 16 h at −80°C, followed by centrifugation at 14,000 rpm for 20 min at 4°C. The supernatant was filtered using biomasher homogenizer tubes (#09-A10-050, OMNI International). Samples were stored at −80°C until use. Protein concentration was determined with the BCA assay (Pierce Biotechnology), followed by normalization. Either 30 μg (for DP2, PPARγ, L-PGDS, Sox-2, COX-2) or 50 μg of protein (for DP1) from each sample were run on 4-12% SDS gels and transferred to nitrocellulose membranes with the iBlot® dry blotting system (Life Technologies) for 7 min. Membranes were blocked with SuperBlock (#37535, ThermoFisher), and hybridized with various primary antibodies followed by HRP-conjugated secondary antibodies (Supplemental Table 10), prior to developing with an enhanced chemiluminescence (ECL) substrate (SuperSignal™ West Pico PLUS, ThermoFisher #34580), and detected on a BX810 autoradiography film (Midwest Scientific). ImageJ software (Rasband, W.S., ImageJ, U. S. National Institutes of Health, Bethesda, Maryland, USA, https://imagej.nih.gov/ij/, 1997-2018) was used for semiquantification by densitometry of the respective bands. Loading controls used were GAPDH, tubulin, or β-actin depending on their molecular weights to avoid overlapping with the other proteins studied.

### 2.8. Cognitive behavior assessment with the passive radial 8-arm maze

This variant of RAM is a passive task that uses positive reinforcement (food) to assess spatial working memory. It is classified as working memory because only short-term memory is used and memory of previous trial baits will not aid the rat in later trials as all baits are used and replenished after each trial. This hippocampal dependent task uses spatial cues in the test room. The maze is divided into eight arms with a bait of food (Ensure® Food Supplement) at the end of each arm. Prior to training, rats were food deprived to 85% of their *ad libitum* body weight and received six shaping trials across two days. For training, the rats were tested four times across two days. The rats begun the training confined to the center of the arena with an opaque covering. Once the opaque covering was removed, the rat was free to start the trial to collect all eight baits. Entrances were recorded after the rat crossed halfway across the arm towards the bait. When the rat returned to a bait that was previously consumed this was deemed as an error. Animals were required to collect all eight baits for the trial to end, and if the trial exceeded 25 min, the trial was not included in the analysis. After each trial, the maze was shifted at 90° and cleaned with a 70% ethanol solution to prevent internal maze cues being used. To prevent the rats using their sense of smell to find baits, the maze room had ample food placed throughout the maze room. Data of all fully completed training trials were analyzed.

### 2.9. Cognitive behavior assessment with the active place avoidance task

This variant of aPAT is an active task that uses negative reinforcement (shock) to access spatial learning. It is classified as reference memory because long-term spatial learning is used as the rats experience repeated trials with a fixed shock quadrant, so referencing previous trials will aid in better performance. The task challenges the rat to avoid a fixed quadrant of the arena as the arena rotates at one revolution per minute. A computer-controlled system was used for aPAT (Bio-Signal Group, Acton, MA). The arena used for this task was enclosed with a transparent plastic wall that was fixed to the arena. An overhead camera (Tracker, Bio-Signal Group) was calibrated to the white hue of the rats and tracked the rat’s movement. This hippocampal dependent task used spatial cues in the test room. The rotating arena forced the rat into the fixed quadrant. After the system detected the rat was in the fixed quadrant for 1.5s, the system delivered a pulse shock of 0.2 milliamperes throughout the arena every 1.5s, giving the rat a foot shock and subsequent foot shocks until it left the fixed quadrant. The rotating arena forced the rat to actively avoid the fixed quadrant, otherwise it would receive a shock. This hippocampal dependent task uses spatial cues to help the rat navigate within the spatial environment. Before training, rats were habituated to the rotating arena for 10 min without a shock. For training, the rats received six 10-min trials with 10-min breaks in their home cage between every trial. To access retention, on the next day the rats received a 10-min trial without a shock zone. The system software recorded data for all trials, and all data was exported to .tbl files and analyzed offline (TrackAnalysis, Bio-Signal Group).

### 2.10. Statistics

All data are represented as the mean ± SEM. Statistical analyses were performed with GraphPad Prism 9 (GraphPad Software, San Diego, CA). All *p* values, SEMs and *t*-statistics are shown on graphs and/or in supplemental tables. Welch’s unpaired one-tailed *t* test was used to compare means between the two groups (WT and Tg-AD) for PG (Fig. 2), IHC (Fig. 3-4, Supplemental Tables 1-7), WB (Fig. 6, Supplemental Table 9), RAM (Fig. 7), and the two groups (TGNT, Tg-AD non-treated, and TGTR, Tg-AD timapiprant-treated males, Fig. 9). Multiple unpaired *t*-test was used for RNAseq (Fig. 5, Supplemental Table 8) for the 21 PG genes with an FDR set to 1% using the two-stage step-up method (Benjamini, Krieger, and Yekutieli). Multi-factor comparisons for aPAT (Fig. 8 and Fig. 9) were performed using a two-way repeated measure analysis of variance (ANOVA), followed by a post hoc (Sidak’s) to access differences across individual training trials or conditions.

**Fig. 2.**
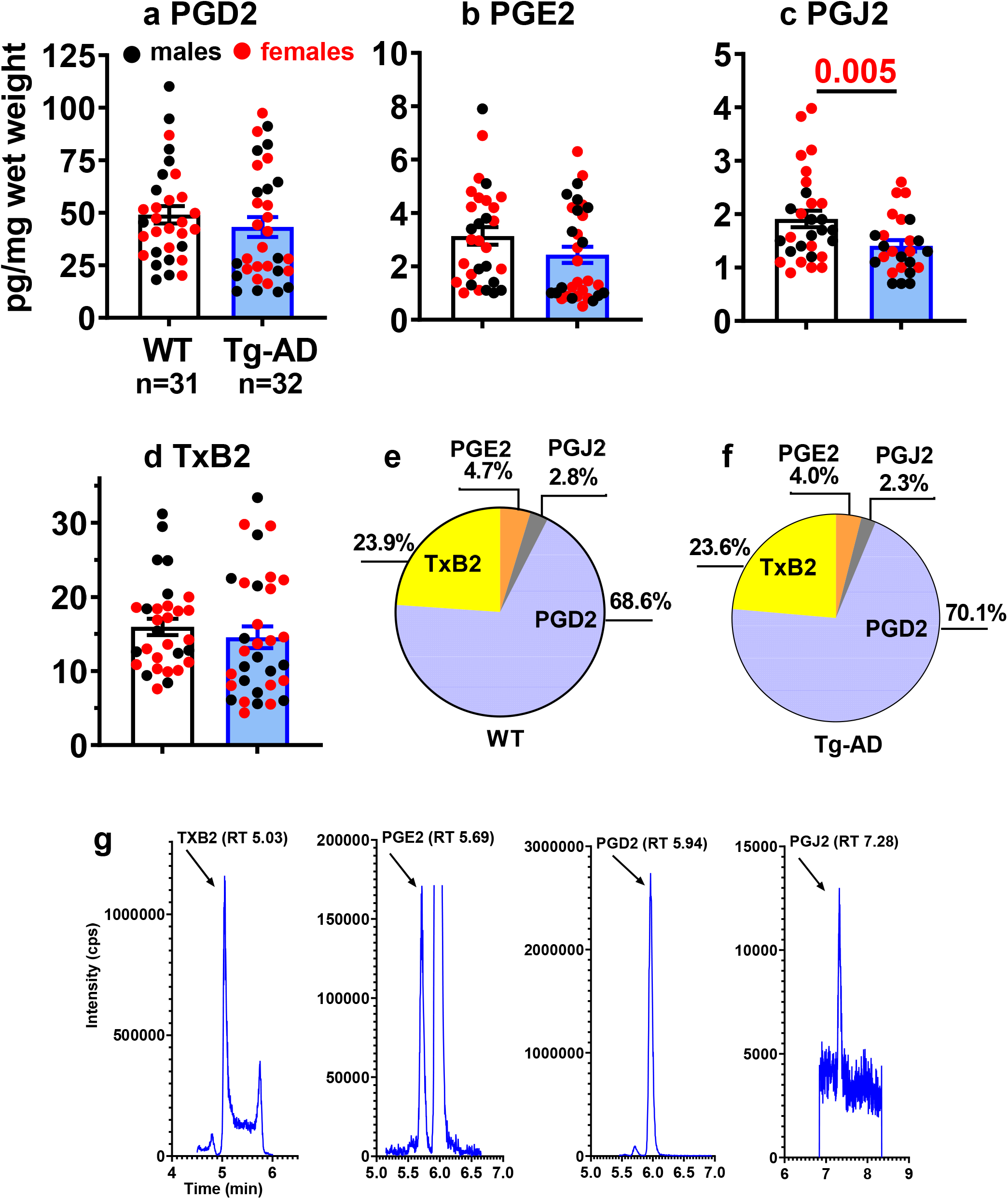
PGD2 is the most abundant PG in the hippocampus of WT and Tg-AD rats Concentrations of prostanoids **a** PGD2, **b** PGE2, **c** PGJ2, and **d** TXB2, measured by LC-MS/MS in whole left hippocampal tissue (combined ventral and dorsal) from 11-month WT (n = 31) and Tg-AD (n = 32) rats. Prostanoids levels were equivalent in Tg-AD and WT littermates, except for PGJ2 that were less (*t* = 2.668, *p* = 0.005). Significance estimated with a two-tailed Welch’s t test. **e-f** Pie graphs represent the proportion of each of the four prostanoids relative to their total sum, in **e** WT and **f** Tg-AD rats. It is clear that PGD2 is the most abundant prostanoid in both WT and Tg-AD rats. **g** Chromatographic profiles depicting the separation of the four prostanoids using LC-MS/MS as explained under materials and methods.

**Fig. 3.**
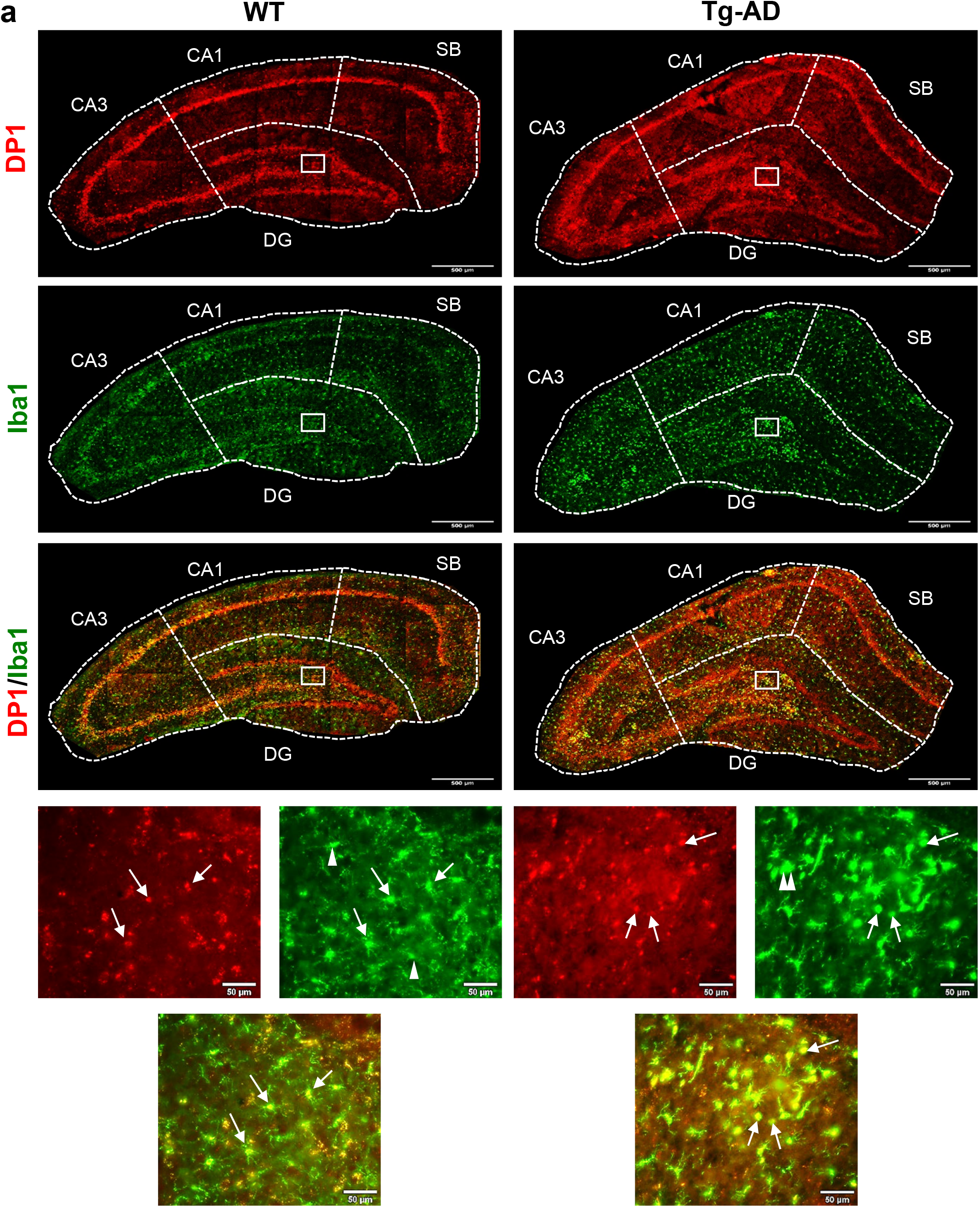
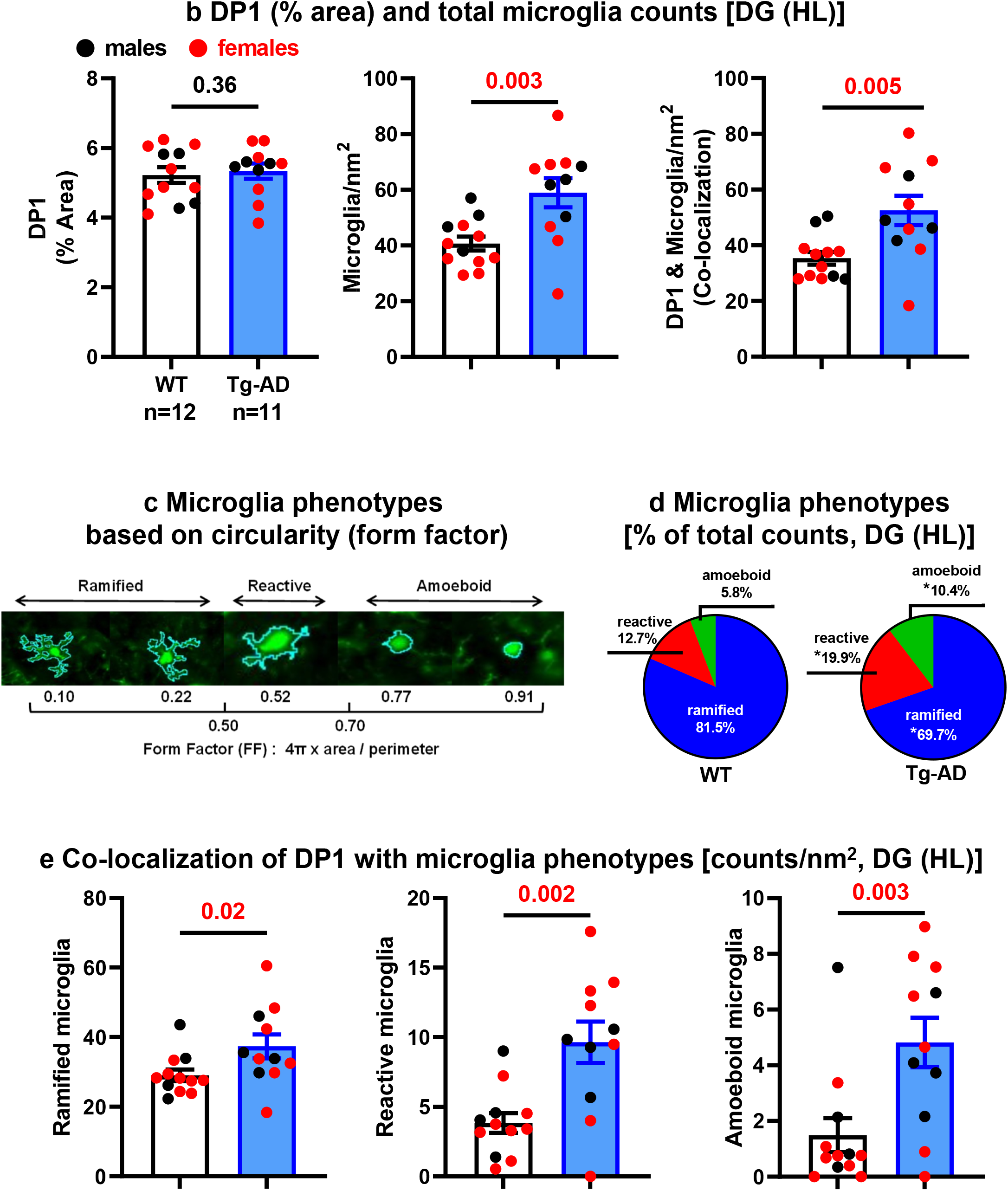
Tg-AD rats have higher levels of microglia and DP1/microglia co-localization in the hippocampus than WT rats **a** DP1 (red), microglia (green, Iba1 antibody), and DP1/microglia co-localization (yellow) IHC analysis of the right dorsal hippocampus of WT (left column, n = 12) and Tg-AD (right column, n = 11). Large panels: 10x magnification, 500 μm scale bars. Small (bottom) panels: 20X magnification of the small white boxes depicted in the larger panels, 50 μm scale bars. White arrows indicate: full, DP1/microglia co-localization; single head, ramified microglia, double head, amoeboid microglia. For DP1 levels, there were no significant differences between WT and Tg-AD rats across all hippocampal regions (**b**, left graph for DG and Supplemental Table 1). Tg-AD rats had significantly more microglia than WT rats, in all hippocampal regions except for CA3 (**b**, middle graph for DG and Supplemental Table 2). Tg-AD rats also had significantly higher levels (1.5 fold) of DP1/microglia co-localization than their WT littermates, only in the DG (HL) region (**b**, right graph for DG only and Supplemental Table 3). **c** The three microglia (Iba1+) phenotypes based on circularity (form factor) as explained under material and methods. **d** Microglia phenotypes as % of total counts at DG (HL). Each pie slice represents the proportion of each phenotype relative to the total sum, in WT (left) and Tg-AD (right) rats. Tg-AD rats had significantly fewer ramified, but more reactive and almost double amoeboid microglia than controls. **e** Co-localization of DP1 with each microglia phenotype in the hippocampal DG (HL) was significantly higher in Tg-AD rats than in WT littermates. Significance (*p* values shown on graphs) estimated by a one-tailed Welch’s *t* test.

**Fig. 4.**
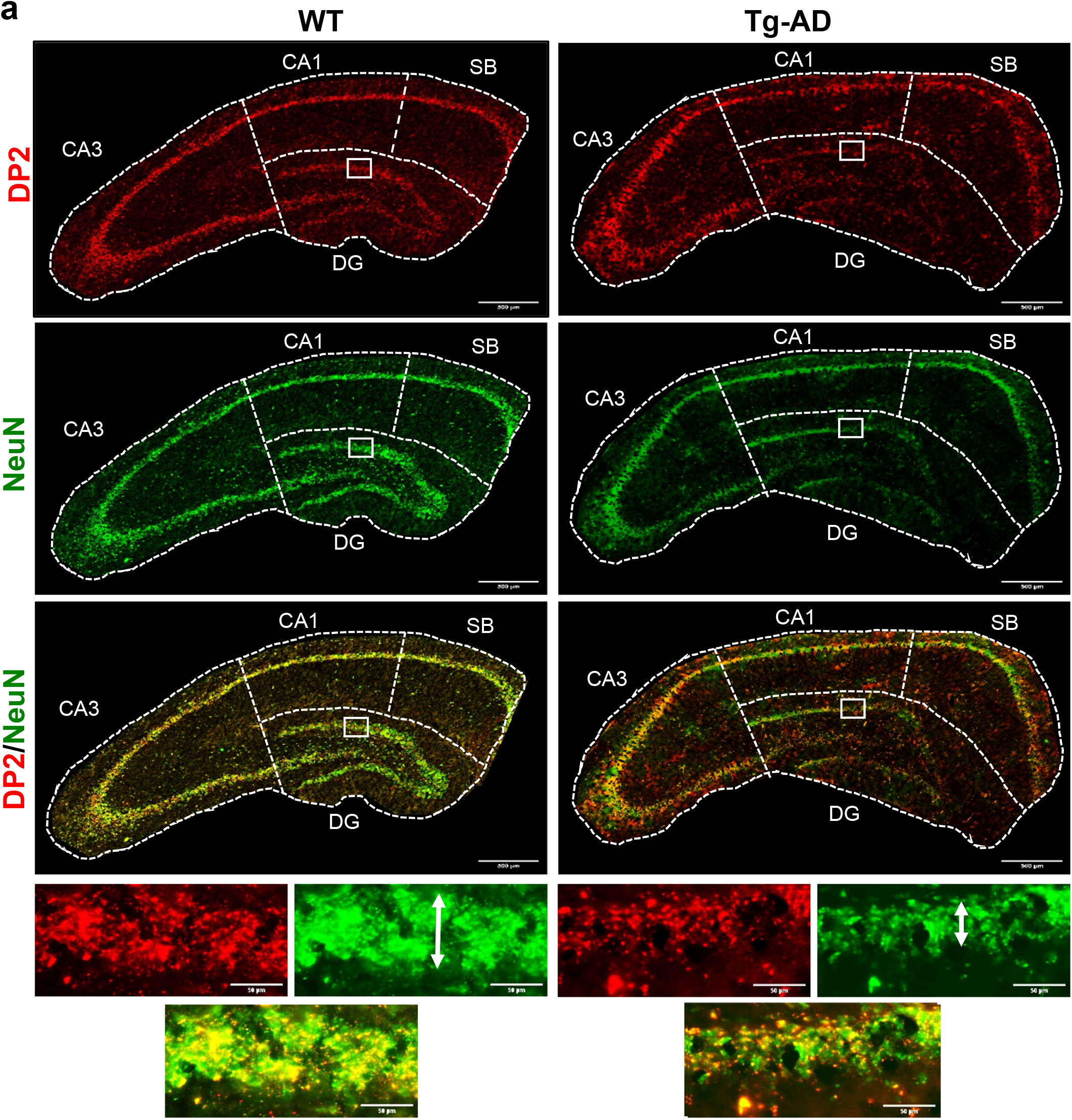
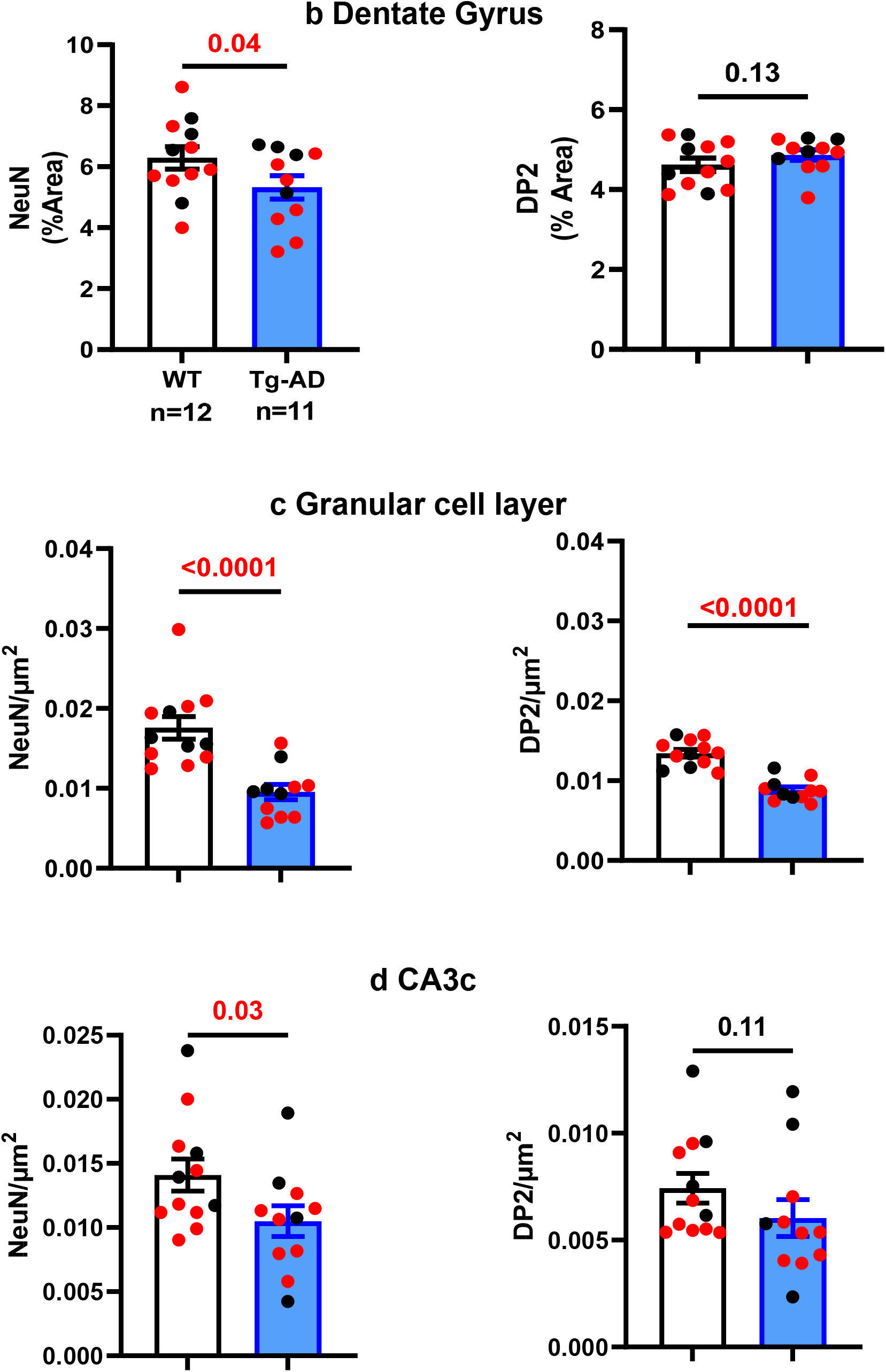
Tg-AD rats display DP2 and neuronal losses in the hippocampus **a** DP2 (red), neurons (green, NeuN antibody), and DP2/neuronal co-localization (yellow) IHC analysis of the right dorsal hippocampus of WT (left column, n = 12) and Tg-AD (right column, n = 11). Large panels: 10x magnification, 500μm scale bars. Small (bottom) panels: 20X magnification of the small white boxes at the GCL depicted in the larger panels, 50μm scale bars. It is clear that the thickness of the GCL is greater in WT than in Tg-AD rats (white double head arrows). (**b – d,** left graphs) Neuronal loss detected only in the DG (**b**), at the GCL (**c**) and CA3c (**d**) of Tg-AD compared to WT rats. Neuronal density analyzed across all other hippocampal regions revealed no changes in Tg-AD compared to WT rats (Supplemental Table 5). (**b – d**, right graphs) DP2 loss detected only at the GCL (**c**) but not at the other hippocampal locations (**b** and **d**, and Supplemental Table 6) of Tg-AD compared to WT rats. Significance (*p* values shown on graphs) estimated by a one-tailed Welch’s t test.

**Fig. 5.**
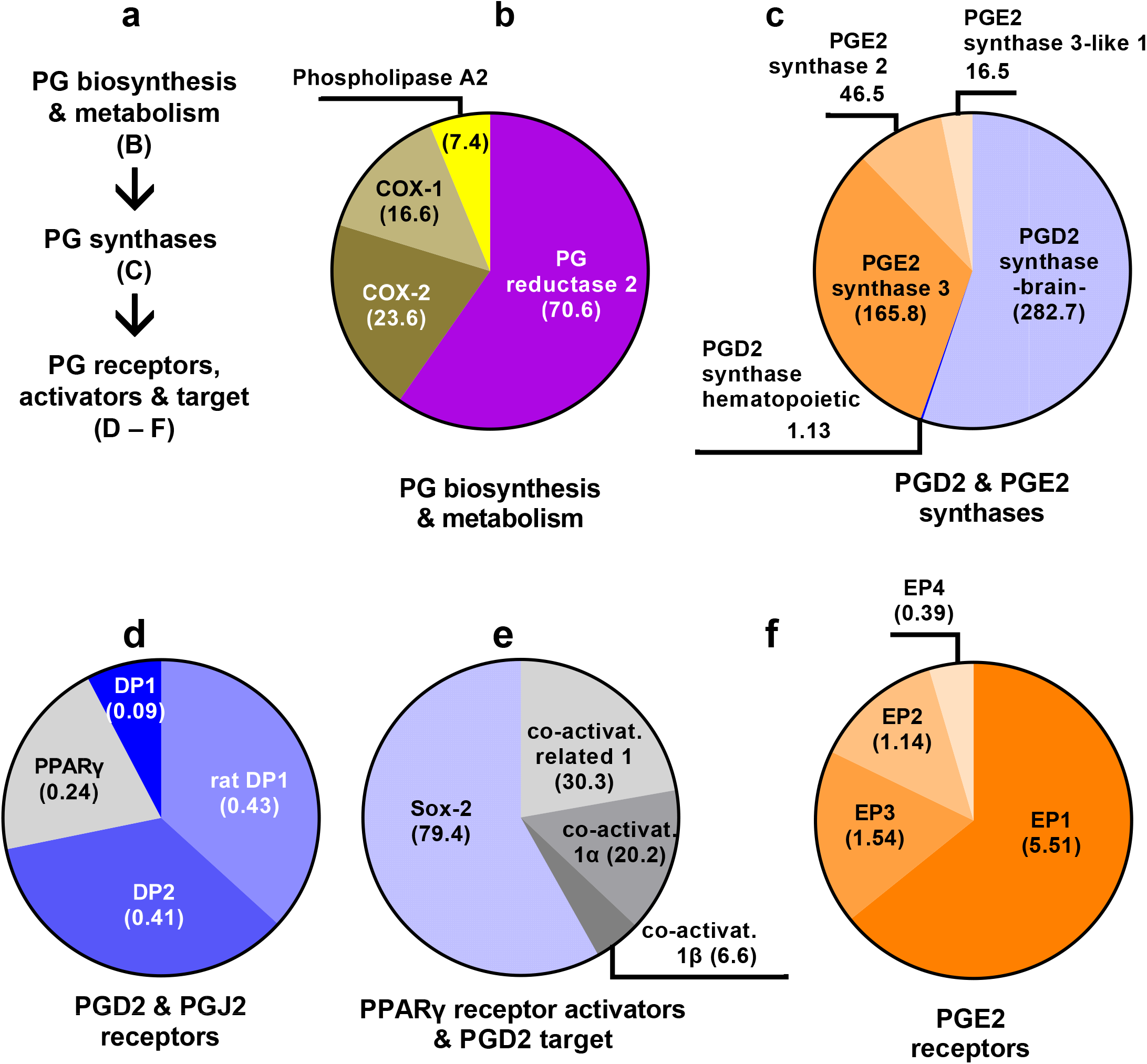
Lipocalin prostaglandin D2 synthase (L-PGDS) mRNA levels are the highest among 21 genes evaluated by RNAseq in the hippocampus of WT and Tg-AD rats **a** The functional groups analyzed within the PG pathway. **b – f** mRNA levels for 21 genes involved in the PGD2 and PGE2 pathways were determined by RNAseq in whole left hippocampal tissue (combined ventral and dorsal) from 11-month WT and Tg-AD male (5 of each genotype) and female (5 of each genotype) rats. The mRNA levels (mean RPMs, WT females only) are displayed as pie graphs, in which each slice is proportional to the total for a specific functional group of genes. The mRNA transcript level was the highest for L-PGDS, the PGD2 synthase in the brain (**c**). Most of the mRNA levels of the 21 genes were not significantly different between Tg-AD rats and WT littermates, except for example the transcription factor Sox-2, which was significantly downregulated in male Tg-AD rats compared to their WT littermates. Additional details are in the text and Supplemental Table 8.

**Fig. 6.**
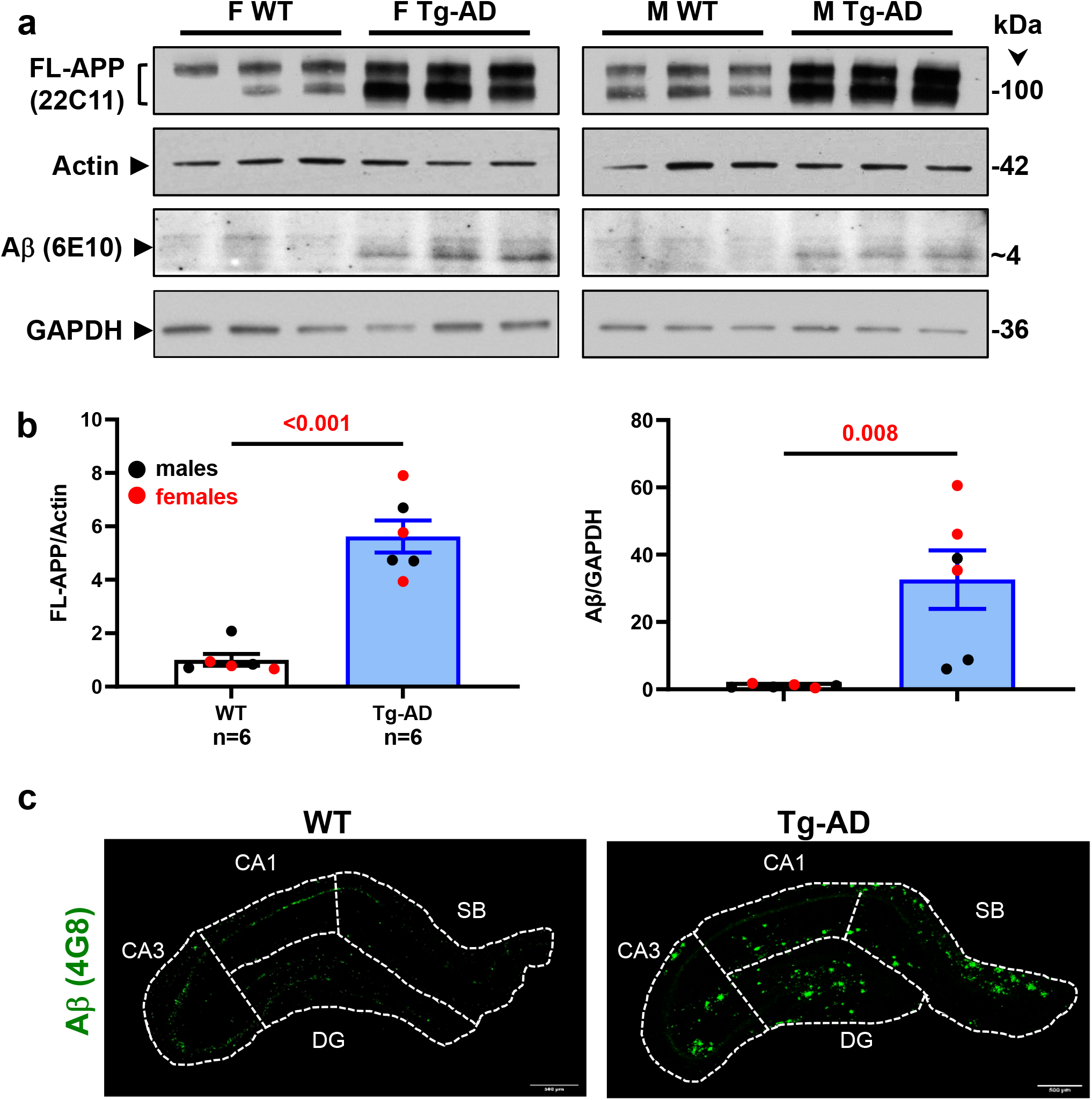
Tg-AD rats show enhanced FL-APP and Aβ peptide levels as well as Aβ plaques in the hippocampus **a** FL-APP (top panels) and Aβ levels (third panels from the top) were assessed by western blot analysis in whole left hippocampal (combined ventral and dorsal) homogenates from 11-month WT and Tg-AD male (M, 3 of each genotype) and female (F, 3 of each genotype) rats. Actin (second panels from the top) and GAPDH (bottom panels) detection served as the respective loading controls. **b** FL-APP and Aβ levels semi-quantified by densitometry. Data represent the percentage of the pixel ratio for FL-APP and Aβ over the respective loading controls for Tg-AD compared to WT (value of one). Values are means ± SEM from 6 rats per genotype (males and females combined). Significance *(p* values shown on graphs) estimated by a one-tailed Welch’s t test. **c** Immunohistochemistry for Aβ plaque load for WT (left panel) and Tg-AD (right panel) rats is shown at 10x magnification, scale bars = 500μm. All Tg-AD rats used in this study exhibited Aβ plaques, but not their WT littermates.

**Fig. 7.**
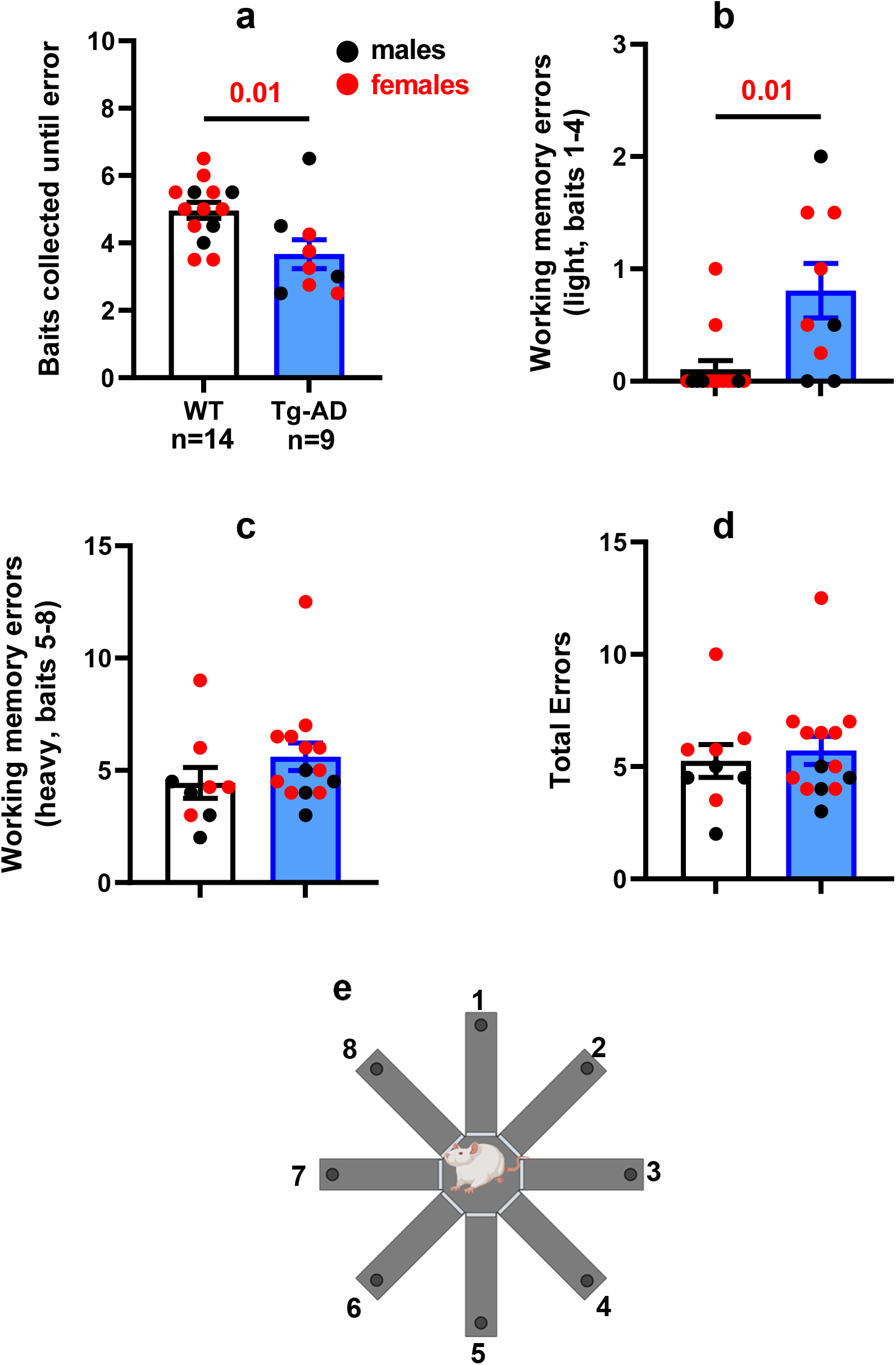
Tg-AD rats have spatial working memory deficits on RAM at 11 months Results show Tg-AD rats (n = 9) commit significantly more errors vs WT rats (n = 14) when analyzed **a** for baits collected until first error, and **b** for working memory errors committed during collection of bait numbers 1-4 (light memory load). **c** No significant differences were observed between conditions with a heavy (challenging) working memory load (collection of baits 5-8), or **d** for total errors committed collecting all 8 baits. Significance estimated with a one-tailed Welch’s t test, and *p* values are shown above bar graphs. **e** RAM with the arms labelled 1 – 8 (created with BioRender.com).

**Fig. 8.**
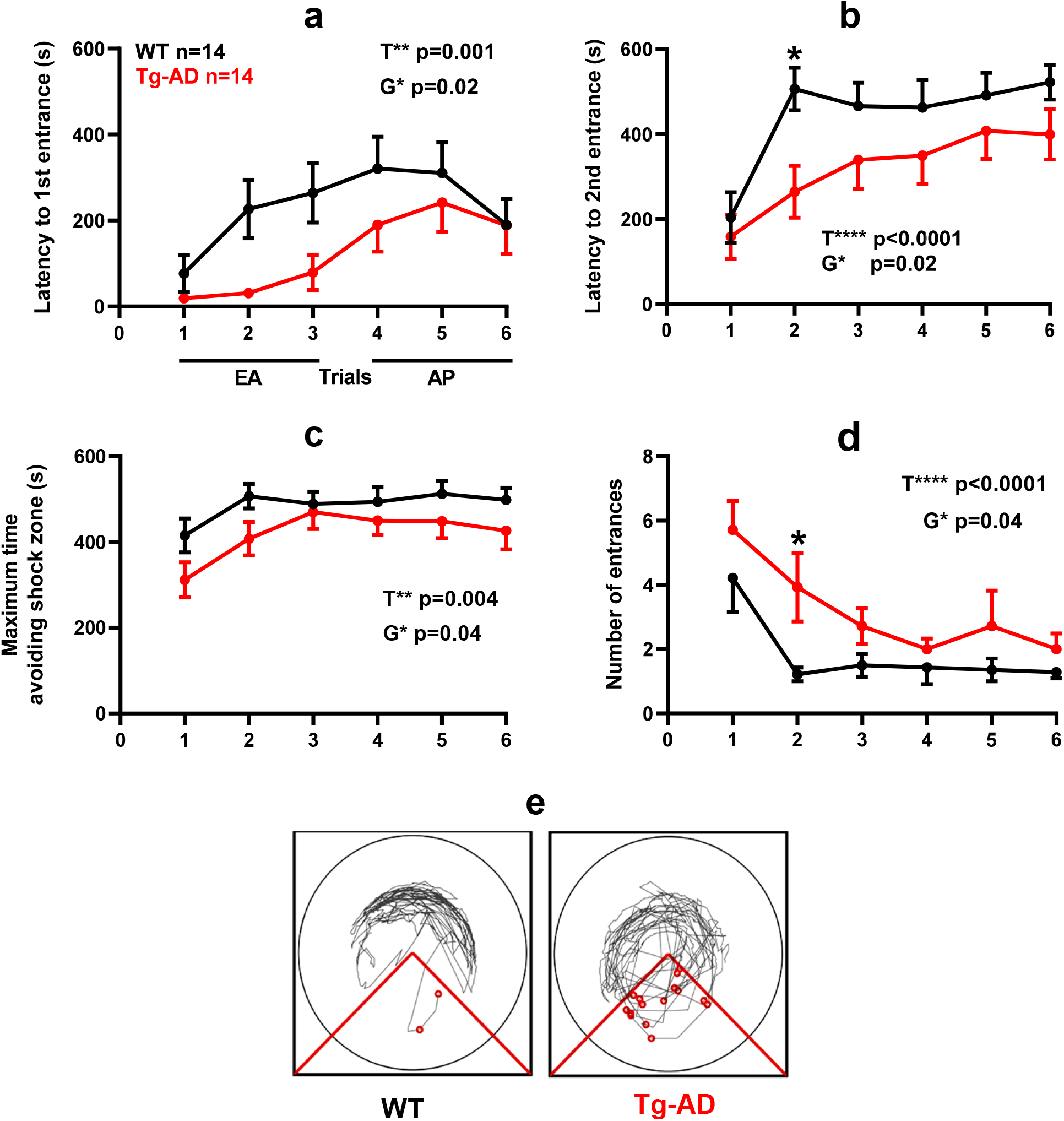
Tg-AD rats exhibit spatial learning deficits on aPAT at 11 months **a-d** All behavioral measures show overall effects by genotype and by trials indicated as *G and *T with corresponding *p* values. **a** Analysis of latency to first entrance shows Tg-AD rats exhibiting significantly shorter latencies. **b** Latency to second entrance shows a significant post-hoc difference on trial 2, \**p* = 0.03. **c** Maximum time to avoid shows Tg-AD rats exhibiting significantly shorter maximum avoidance latencies. **d** Number of entrances shows a significant post-hoc difference on trial 2, \**p* = 0.03. Representative track tracings for trial 2 shown for WT and Tg-AD female rats. Significance estimated by a two-way repeated measure ANOVA with Sidak’s post hoc for multiple comparisons.

**Fig. 9.**
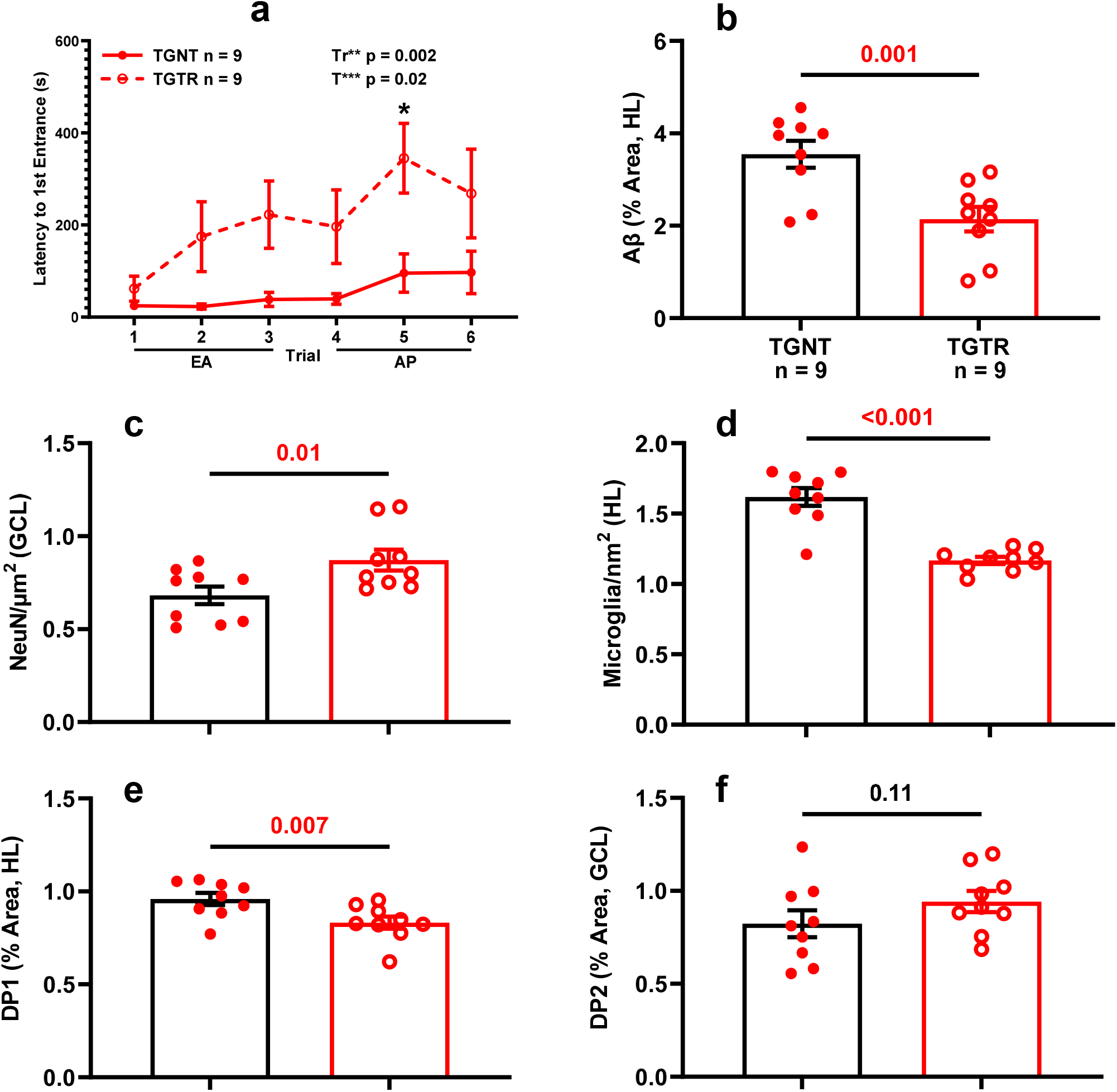
Timapiprant mitigates AD pathology. Tg-AD timapiprant-treated (TGTR, n = 9) compared to Tg-AD nontreated (TGNT, n = 9) male rats perform significantly better in latency to first entrance during training (**a**, *p* = 0.002) with a significant post-hoc difference on trial 5, experience lower plaque burden (**b**, *p* = 0.001), higher neuronal levels (**c**, *p* = 0.01), and lower microglia levels (**d**, *p <* 0.001) in the DG hilar subregion of the hippocampus. DP1 and DP2 receptor levels were decreased (**e**, *p* = 0.007) and unchanged (**f**, *p* = 0.11), respectively, in the same hippocampal subregion of TGTR vs TGNT male rats. Unpaired one-tail *t*-tests with Welch’s corrections were used for statistical analysis. EA - early acquisition; AP – asymptotic performance; T**, training effect; Tr*** = timapiprant-treatment effect; (\**p* < 0.05; ** *p* < 0.01; *** *p* < 0.001.

## 3. Results

### 3.1. PGD2 is the most abundant PG in the hippocampus of WT and Tg-AD rats

Rat hippocampal tissue from 11-month WT (n = 31) and Tg-AD (n = 32) rats was analyzed to determine PGD2, PGE2, PGJ2 and thromboxane B2 (TxB2) concentrations. Quantitative levels of the four prostanoids ranged from 0.7 to 110.2 pg/mg wet tissue. The levels measured in the order of abundance were PGD2, TxB2, PGE2, and PGJ2, at 49.1 ± 4.1, 17.0 ± 1.5, 3.4 ± 0.4, and 2.0 ± 0.2 pg/mg wet tissue for WT rats (Fig. 2a-d). Prostanoid levels were similar in Tg-AD and WT littermate. Quantitative amounts of the four prostanoids in Tg-AD rats measured in the order of abundance were PGD2, TxB2, PGE2, and PGJ2, at 43.2 ± 4.7, 14.6 ± 1.5, 2.4 ± 0.3, and 1.4 ± 0.1 pg/mg wet tissue (Fig. 2a-d). In seven Tg-AD rats, PGJ2 was not detectable. The latter is produced from PGD2 by non-enzymatic dehydration and its formation in vivo remains controversial [9, 15]. All of these values are in accordance with those previously reported for Sprague-Dawley male rat brain cortical tissue at postnatal day 16–18, measured by quantitative UPLC–MS/MS [57, 58]. Under normal conditions, quantitative amounts of PGD2, PGJ2, and PGE2, measured in the order of abundance were 123.7, 12.3, and 4.5 pg/mg wet tissue, or 351, 36.9, and 12.8 pmol/g wet tissue [57, 58]. The differences between the latter study and ours can be accounted for by the prostaglandin levels being quantified in different rat strains, at different ages and in different brain regions.

Out of the four prostanoids measured, PGD2 was by far the most abundant in the hippocampal tissue, as reflected in the pie graphs shown in Fig. 2e-f. These graphs represent the proportion of each of the four prostanoids relative to their total sum. For example, it is clear that PGD2 levels are 14.5-fold and 17.5-fold higher than PGE2 in WT and Tg-AD rats, respectively (Fig. 2e-f). PGD2 levels represent 68.8% and 70.1% relative to total, while PGE2 levels represent 4.7% and 4.0% relative to total in WT and Tg-AD rats, respectively.

The specific chromatographic profiles of calibration standards for each prostanoid is depicted in Fig. 2g, showing that the four prostanoids derived from arachidonic acid can be quantified reliably in rat hippocampal tissue using LC-MS/MS analysis. Under our experimental conditions, the elution sequence was identified as TxB2, PGE2, PGD2, and PGJ2.

### 3.2. Tg-AD rats have enhanced microglia and DP1/microglia co-localization levels in the hippocampus

We assessed DP1 and microglia levels in the hippocampus of WT and Tg-AD rats at 11 months of age [Fig. 3a: DP1, red; microglia, green; DP1/microglia co-localization, yellowish (indicated by single white arrows)]. It is clear that DP1 is detected in the four discrete hippocampal regions (SB, CA1, CA3, and DG) in WT and Tg-AD rats (Fig. 3a). For DP1 levels, there were no significant differences between WT and Tg-AD rats, considering the four hippocampal regions individually (Fig. 3b, left graph for DG only and Supplemental Table 1 for all). A different situation was observed for microglia, as Tg-AD rats had significantly more microglia than WT rats, in all hippocampal regions except for CA3 (Supplemental Table 2). For example, Tg-AD rats had more microglia (1.4 fold, *t* = 3.15, *p* = 0.003) in the DG hilar (HL) subregion than their WT littermates [Fig. 3b, middle graph for DG (HL) only)].

It is evident that DP1 is co-localized with microglia in all four hippocampal regions (Fig. 3a), shown at higher magnification for the DG (HL) (Fig. 3a, bottom panels, indicated by single white arrows). Tg-AD rats had significantly higher levels (1.5 fold, *t* = 2.99, *p* = 0.005) of DP1/microglia co-localization than their WT littermates, only in the DG (HL) region [Fig. 3b, right graph for DG (HL) only, and Supplemental Table 3 for all].

Microglia have a remarkable variety of morphologies associated with their specific functions, and can be divided into three phenotypes according to their cell body circularity: ramified, reactive and amoeboid [31] (Fig. 3c). Notably, when compared to WT controls, Tg-AD rats showed a shift from a neuroprotective state typical of ramified microglia, to more of a neurotoxic and overactive state attributable to amoeboid microglia. In the hippocampal DG (HL), there are significant less ramified microglia (14.5 % less, *t* = 2.32, *p* = 0.02) with a concomitant increase in reactive (1.6 fold more, *t* = 2.52, *p* = 0.01) and amoeboid (1.8 fold more, *t* = 1.89, *p* = 0.04) microglia in Tg-AD rats compared with WT controls (Fig. 3d).

Co-localization of DP1 with each microglia phenotype in the hippocampal DG (HL) was significantly higher in Tg-AD rats than in WT littermates (Fig. 3e). Accordingly, compared to WT controls, the Tg-AD rats had 1.3 fold (*t* = 2.19, *p* = 0.02), 2.5 fold (*t* = 3.53, *p* = 0.002), and 3.2 fold (*t* = 3.08, *p* = 0.003) higher DP1 co-localization with ramified, reactive and amoeboid microglia, respectively.

No significant differences between WT and Tg-AD rats were detected for astrocyte levels by IHC analysis at 11 months of age in all hippocampal regions (GFAP levels, Supplemental Table 4).

### 3.3. Tg-AD rats display DP2 receptor and neuronal losses in the hippocampus

Neuronal density across hippocampal regions (SB, CA1, and CA3 pyramidal cell layers, and DG granular cell layer) were assessed with NeuN (green) to quantify mature neurons (Fig. 4a-d and Supplemental Fig. 1). Similar to what was reported in the original study for Tg-AD rats at 16 and 26 months of age [13], we observed a significant neuronal loss (NeuN signal) though earlier, at 11 months of age. We detected neuronal loss in Tg-AD compared to WT rats, only in the granular cell layer (GCL) and CA3c pyramidal cell layer of DG (44.2% less, t = 4.75, *p* < 0.0001 for GC, and 21.4% less, *t* = 2.07, p = 0.03 for CA3c) (Fig. 4a-d, left graphs). It is clear that the thickness of the GCL is greater in WT than in Tg-AD rats, shown in higher magnification (Fig. 4a, bottom panels indicated by white double head arrows). Neuronal levels analyzed across all other hippocampal regions revealed no changes in Tg-AD compared to WT rats (Supplemental Table 5).

A similar trend was detected for DP2 receptor levels (red) at the GCL only. Tg-AD rats exhibited significantly fewer DP2 receptors in GCL than the WT controls (34.4% less, *t* = 7.25, *p* < 0.0001 for GCL) (Fig. 4a and c, and Supplemental Table 6). In all other hippocampal regions there were no differences in DP2 levels between WT and TG-AD rats, except in the CA1 region, where DP2 levels were 1.3 fold higher in Tg-AD than in WT controls (*t* = 3.36, *p* = 0.002 and Supplemental Table 6). The observed DP2 increase in the CA1 region of Tg-AD rats is likely due to the presence of Aβ plaques, which disorder the area aspect.

It is clear that at least 50% of DP2 is co-localized with neurons, as shown in Fig. 4a (yellow). For this reason, DP2 receptor and neuronal co-localization was not significantly different between WT and Tg-AD rats at all hippocampal regions. This finding supports that at least 50% of NeuN and DP2 signals are co-localized, and that their decrease in Tg-AD compared to WT, follows the same trend (Supplemental Table 7).

### 3.4. Lipocalin prostaglandin D2 synthase (L-PGDS) mRNA levels are the highest among 33 genes evaluated by RNAseq in the hippocampus of WT and Tg-AD rats

We assessed the mRNA levels for 33 genes involved in the PGD2 and PGE2 pathways in hippocampal tissue from WT and Tg-AD male (5 of each genotype) and female (5 of each genotype) rats. The RNAseq analysis reports output measures as reads per million (RPM), as well as false discovery rate (FDR) and *p* values (Supplemental Table 8). In addition, the mRNA levels (mean RPMs) for 21 of those genes are displayed as pie graphs, in which a slice of each pie is proportional to the total for a specific functional group of genes (Fig. 5). The functional groups within the PG pathway are depicted in Fig. 5a. The numbers shown on the pie graphs (Fig. 5b-f) represent the RPMs from WT females only, as there were no significant genotype (WT vs Tg-AD) or sex (male vs female) differences in the expression levels for most of these genes (Supplemental Table 8).

Overall, the data reveal that mRNA transcript levels were the highest for L-PGDS, the PGD2 synthase in the brain (RPM = 282.7, Fig. 5c). This is consistent with PGD2 being the most abundant prostanoid of the four that we measured in hippocampal tissue (as much as 70% of the total, Fig. 2e and f). Hematopoietic-PGDS (H-PGDS), the other PGD2 synthase that is mainly detected in microglia [42], was minimally expressed (RPM = 1.13, Fig. 5c). There were three PGE2 synthases detected by RNAseq, and their mean RPMs were in descending order, 165.8 (PGES-3), 46.5 (PGES-2), and 16.5 (PGES-3-like1) (Fig. 5c).

Evaluation of four genes involved in PG biosynthesis and metabolism (Fig. 5b), showed that prostaglandin reductase-2 (pTGR-2), which metabolizes PGs, exhibited the highest expression (RPM = 70.6). The remaining three genes followed in descending order, COX-2 (RPM = 23.6), COX-1 (RPM = 16.6) and phospholipase A2 (RPM = 7.4).

In the rat hippocampal tissue, the mRNA levels (RPM) for PGD2 receptors were, in descending order, as follows (Fig. 5d): DP1 (rat, orthologous to human DP1, 0.43), DP2 (0.41), and DP1 (0.09). The rat genome has two DP1 copies (genes: PTGDR, ID: 63889 and PTGDRL, ID: 498475). The protein alignments are highly similar (354/357 residues, 99% homology, NCBI groups the two in an identical protein group), differing only on their location on chromosome 15. RPMs for PPARγ, a putative PGJ2 receptor, were = 0.24. Notably, PPARγ activators were expressed at higher levels than the receptor itself (Fig. 5e): co-activator related 1 (30.3), co-activator 1α (20.2), and co-activator 1β (6.6). The receptors for PGE2 showed the highest RPM levels (Fig. 5f) listed in descending order: EP1 (5.51), EP3 (1.54), EP2 (1.14), and EP4 (0.39).

The SRY-box transcription factor 2 (Sox-2) gene is a transcription factor best known as a reprogramming factor necessary for generating induced pluripotent stem cells [61]. Sox-2 is also required for proliferation and differentiation of oligodendrocytes during postnatal brain myelination and CNS remyelination [71]. In addition, Sox-2 is a negative regulator of myelination by Schwann cells [20], and its levels are controlled by L-PGDS in PNS injured nerves [22]. We found that Sox-2 expression levels in the hippocampal tissue were quite high (79.4 RPM, Fig. 5e), being the third highest expressed gene in our list, after L-PGDS and PGES-3 (Supplemental Table 8). Moreover, Sox-2 was significantly downregulated in male Tg-AD rats compared to their WT littermates (22.2 % less, *p* = 0.011, Supplemental Table 8). The significance of these data will be addressed in the discussion.

RNAseq pie graphs for WT and Tg-AD male and female rats are shown in Supplemental Fig. 2-6. In addition, western blot analyses for six proteins involved in the PGD2 pathway, i.e. receptors DP1, DP2, and PPARγ, the synthase L-PGDS, as well as COX-2 and Sox-2, are shown in Supplemental Fig.7. The data show no changes in the levels of these protein in hippocampal tissue from WT and Tg-AD male (n = 3 for each genotype) and female (n = 3 for each genotype) rats (Supplemental Table 9). The whole image of the western blots for DP1, DP2 and L-PGDS is shown in Supplemental Fig. 8.

### 3.5. Tg-AD rats show enhanced FL-APP and Aβ peptide levels as well as Aβ plaques in the hippocampus

The original study by Cohen et al reported that Tg-AD rats express 2.6-fold higher levels of human full-length APP than their WT littermates, assessed by western blot analysis of the brain [13]. Full length APP (FL-APP) was detected with the mouse monoclonal antibody 22C11, which reacts with human and rat, as well as other species (manufacturer’s specifications). In our studies using the same antibody, it is clear that the levels of FL-APP are 5.6 fold higher in the hippocampal tissue of Tg-AD than WT rats (Fig. 6a, top panels labeled with FL-APP, and Fig. 6b, left graph, combined males and females, *t* = 7.23, *p* < 0.001). This trend was observed in males (n = 3 for each genotype) and females (n = 3 for each genotype), and the values were normalized for actin (Fig. 6a, second panels). The 2.2-fold higher levels of FL-APP detected in our analysis compared to Cohen et al [13], could be explained by our studies evaluating hippocampal tissue while whole brain tissue was used in the Cohen et al studies [13].

We also assessed Aβ levels in the same samples of rat hippocampal tissue with the mouse monoclonal antibody 6E10, which has a 3-fold higher affinity for human APP and Aβ compared to rat (manufacturer’s specifications). Aβ peptides were detected in male and female Tg-AD rats but not in the WT littermates, as shown in Fig. 6a [third panels labeled with Aβ (6E10)], and semi-quantified in Fig. 6b (right graph, combined males and females, *t* = 3.62, *p* = 0.008). The whole images of the western blots for FL-APP and Aβ are shown in Supplemental Fig. 9.

The presence of Aβ plaques in all 11-month Tg-AD rats was confirmed by IHC analysis with the mouse monoclonal antibody 4G8, as shown for a female rat in Fig. 6c, right panel. A WT female rat is included for comparison (Fig. 6c, left panel).

### 3.6. Tg-AD rats exhibit impaired spatial learning and memory

In the original studies with the Tg-AD rats [13], cognitive behavior was assessed at 6, 15 and 24 months of age. Most of the significant changes between WT and Tg-AD rats were detected at 15 and 24 months, but not at 6 months of age [13]. To shorten the experimental time line, we evaluated the WT and Tg-AD rats at an earlier age, i.e. at 11 months. We evaluated cognitive impairment with two hippocampal-dependent tasks to measure short-term learning/memory and navigation: 8-arm radial arm maze (RAM), which is a passive behavioral task, and the active-place avoidance task (aPAT). Since in the original studies no sex differences were reported [13], males and females were combined for our analyses.

In our RAM studies, we wanted to assess total errors made, baits collected until error, and working memory. Two forms of working memory were assessed, a light working memory load for baits 1-4, and a heavy (challenging) working memory load for baits 5-8 (Fig. 7e). For RAM the rat groups were WT (10 females, 4 males) and Tg-AD (5 females and 4 males). We found that Tg-AD rats had a behavioral deficit in outputs for baits collected until error (Fig. 7a, *t* = 2.65, *p* = 0.01) and in light working memory load (Fig. 7b, *t* = 2.75, *p* = 0.01). We found no differences in the more difficult measures such as heavy working memory load (Fig. 7c; *t* = 1.26, *p* = 0.11) and total errors (Fig. 7d, *t* = 0.48, *p* = 0.32). These findings show a working memory impairment in the early and less challenging part of the task for the Tg-AD rats compared to controls.

In the aPAT analysis, we used a separate cohort of rats. In aPAT, the rat groups were WT (7 females, 7 males) and Tg-AD (8 females, 6 males). We found a significant deficit in spatial reference memory during training for the Tg-AD rats in all of the reported measures: latency to 1^st^ entrance (Fig. 8a, F_(1,26)_ = 5.73, *p* = 0.02), latency to 2^nd^ entrance (Fig. 8b, F_(1,26)_=5.70, *p* = 0.02), maximum time avoidance (Fig. 8c, F_(1,26)_=4.74, *p* = 0.04) and entrances (Fig. 8d, F_(1,26)_ = 4.78, *p* = 0.04). Significant post hoc differences were observed at trial 2 during the early acquisition (EA) phase in latency to 2^nd^ entrance (Fig. 8b, *t* = 3.05, *p* = 0.03), and in the number of entrances (Fig. 8d, *t* = 2.81, *p* = 0.03).

### 3.7. Timapiprant improves spatial learning and mitigates plaque burden, neuronal loss and microgliosis

We used the hippocampal-dependent active place avoidance task to assess short-term working memory performance on 11-month old Tg-AD non-treated (TGNT) and Tg-AD timapiprant-treated (TGTR) male rats, and the equivalent WT male littermates. The measurement latency to first entrance into the shock zone for TGTR vs TGNT revealed an overall significant effect of treatment (Fig. 9a, F_(1, 16)_ = 13.87, *p* = 0.002) and of training (Fig. 9a, F_(5,80)_ = 2.93, *p* = 0.02). Significant post hoc differences were observed at trial 5 during the asymptotic performance (AP) phase in latency to 1^st^ entrance (Fig. 9a, *t* = 3.15, *p* = 0.01) between TGTR and TGNT rats.No differences were detected between WTNT and WTTR males (F_(1, 16)_ = 1.04, *p* = 0.32), nor between WTNT and TGTR (F_(1, 16)_ = 0.88, *p* = 0.36), showing the beneficial effects of timapiprant-treatment only under pathological conditions. Graphed data (9c-9e) represent a ratio of the Tg-AD rats over their WT controls.

We compared Aβ plaque burden in the hippocampal DG hilar subregion between TGTR and TGNT rats and found that timapiprant significantly mitigated Aβ plaque load (Fig. 9b, *t* = 3.55, *p* = 0.001). Similarly, timapiprant alleviated neuronal loss in the GCL subregion (Fig. 9c, *t* = 2.56, *p* = 0.01) and microgliosis in the hilar subregion (Fig. 9d, *p* < 0.001) for TGTR compared to TGNT male rats. DP1 receptor levels were decreased in the DG hilar subregion of TGTR compared to TGNT male rats (Fig. 9e, *t* = 2.78, *p* = 0.007), but not those of the DP2 receptor in the GCL (Fig. 9f, *t* = 1.29, *p* = 0.11).

## 4. Discussion

There is still much to learn about the profile and role of PGs in AD pathology. We focused on the PGD2 pathway, because it is the most abundant PG in the brain, and its contribution to AD merits more attention. Much more is known about the relation between the PGE2 pathway and AD [67]. Characterizing key elements of the PGD2 pathway in AD could serve as potential biomarkers and/or therapeutic targets for treating this devastating disease.

We investigated the PGD2 pathway in the Tg-AD rats at 11 months of age, because it is midway between the ages at which these rats present mild (at 6 months of age) and robust (at 16 months of age) AD pathology, as reported in the original study [13]. Understanding pre- and/or early-stages of AD is paramount, as treating AD at these stages would be the best approach to preventing severe progression [59]. We established that at 11 months of age the Tg-AD rats exhibit impaired hippocampal-dependent spatial learning and memory, as well as molecular markers of AD, such as amyloid plaques, microglial activation, neuronal loss, and early signs of tau-PHF, the latter reported in our submitted manuscript [46].

Interestingly, we found that neuronal loss at 11 months of age was specific to the DG and its subregions GCL and CA3c in the hippocampal tissue. The DG is known to be vulnerable to aging and to be affected in the early stages of AD [62]. In fact it is reported that in AD the GCL of the DG has impaired firing [49]. When the GCL is impaired, the ability to identify/discriminate environmental cues during memory formation is greatly diminished in spatial learning/memory [33]. These data support our findings that at 11 months of age, Tg-AD rats exhibit a significant impairment in two separate hippocampal-dependent behavioral tasks where the use of environmental cues is necessary to evaluate learning/memory.

The relative abundance of the four prostanoids that we measured in hippocampal tissue was, in descending order, PGD2, TxB2, PGE2, and PGJ2. PGD2 was by far the most abundant at ~46.2pg/mg wet tissue, as it was ~ 3-fold higher than TxB2, ~16-fold higher than PGE2, and ~28-fold higher than PGJ2, all reported as an average between WT and Tg-AD rats since there was no significant difference between the two genotypes. Other studies confirm our finding that PGD2 is the most abundant PG in the brain, including in human brains [24, 45, 52, 57].

PGD2 levels were equivalent in Tg-AD and WT rats. In contrast, one study reported that PGD2 levels were significantly higher in post-mortem frontal cerebral cortex tissue from AD patients compared to age matched controls [27]. Moreover, others demonstrated that PGD2 levels increase significantly by as much as 6-fold in the hippocampal and/or cerebral cortical tissue of male Sprague-Dawley rats following traumatic brain injury [32] or brain ischemia [37, 38, 57]. Several factors could explain why the levels of PGD2 were equivalent in the hippocampus of 11-month Tg-AD and WT rats. Firstly, the studies with the human AD cases [27] measured PGD2 levels in a different brain area, i.e. the cerebral cortex. Secondly, PG levels were assessed in the human cerebral cortices upon a 30-min incubation of the tissue at 37°C, thus assessing PGD2 produced *de novo* during those 30-min. Under these conditions, PGD2 levels were significantly (~2-fold) higher in the AD cases than in controls. In contrast, there were no marked differences for PGE2 between the AD cases and controls [27]. Thirdly, the short half-life of PGD2 could explain the discrepancy between our studies and those involving different forms of rat brain injury. The half-life of PGD2 in mice was estimated to be 1.6 min in the brain and 1.5 min in the blood [60]. Therefore, the increase in the levels of PGD2 under a chronic condition, such as in AD addressed with our studies, could be harder to detect than soon after brain injury such as that induced by traumatic brain injury [32] or brain ischemia [37, 38, 57]. Although the exact cause of the PGD2 increase is unclear, PGD2 production could be accelerated to compensate for neuronal damage, and possibly enhance neuronal activity in the injured brain, as suggested by [27]. Clearly, further investigation into this matter is needed.

The biologic actions of PGD2 are elicited through binding to its receptors DP1 and DP2 on specific cell types. In the brain, DP1 was detected in microglia [43], astrocytes [43], and neurons [35]. Moreover, DP1 was specifically localized in microglia and reactive astrocytes associated with senile plaques in the cerebral cortex of AD patients and of Tg2576 mice, a model of AD [42]. In our studies with 11-month Tg-AD and WT rats, we detected changes in hippocampal microglial numbers but not in astrocytes, thus we investigated DP1 distribution among the three microglia phenotypes, i.e. ramified, reactive and amoeboid. Notably, in the hippocampal hilar subregion, Tg-AD rats had significantly fewer ramified, but more reactive and almost double amoeboid microglia than controls. Thus, the Tg-AD rats at 11 months of age, exhibited a shift from a neuroprotective state typical of ramified microglia, to more of a neurotoxic and overactive state attributable to amoeboid microglia. This was expected as the latter state is associated with neurodegeneration [11].

We established that DP1/microglia co-localization at the hippocampal hilar subregion increased the most (3.2-fold) in amoeboid microglia of Tg-AD rats compared to controls. Enhanced DP1/amoeboid microglia colocalization could be considered an early marker of neurodegeneration. In fact, microglial overactivation and recruitment are induced by Aβ, leading to microglia clustering around Aβ aggregates at an early stage prior to neuropil damage in AD patients [11]. Furthermore, microglia-mediated neurotoxicity manifested by the production of pro-inflammatory cytokines such as TNFα and IL-1β, reactive oxygen/nitrogen species, and chemokines [4], tends to be progressive potentially contributing to the progressive nature of AD [11].

Whether the increase in DP1/amoeboid microglia co-localization contributes to the neurodegenerative process or is a compensatory mechanism, remains to be established. Both DP1 agonists and antagonists can be protective in the brain and/or spinal cord, depending on the type of injury (Fig. 10). On the one hand, DP1 agonists such as BW245C protect against glutamate toxicity and ischemic stroke induced in rodents [3, 35]. The benefits of DP1 activation are mediated by increased cAMP synthesis that is instrumental in converting pro-inflammatory neurotoxic microglia towards a tissue reparative anti-inflammatory phenotype [23]. Among other effects, DP1 activation facilitates vasodilation, thus protecting the brain from ischemic stroke caused by brain blood vessels becoming clogged [2]. DP1 activation also regulates sleep by stimulating adenosine formation and subsequently activating the adenosine receptor A2A [5]. Studies with mice showed that sleep drives Aβ clearance from the adult brain [69]. Both ischemic stroke [65] and sleep dysregulation [29] facilitate the progression of AD pathology. On the other hand, by limiting bleeding in mice, DP1 antagonists such as laropiprant (MK-0524) protect against hemorrhagic stroke caused by brain bleeding that affects its function [4]. Moreover, DP1 genetic ablation mitigated disease symptoms developed by a mouse model of amyotrophic lateral sclerosis (ALS) [16]. DP1 inhibition mitigates the increase in activated/amoeboid microglia associated with both hemorrhagic stroke and ALS [4, 16]. In conclusion, modulating DP1 function is a promising therapeutic strategy applicable to different types of brain conditions and injuries related to AD.

**Fig. 10.**
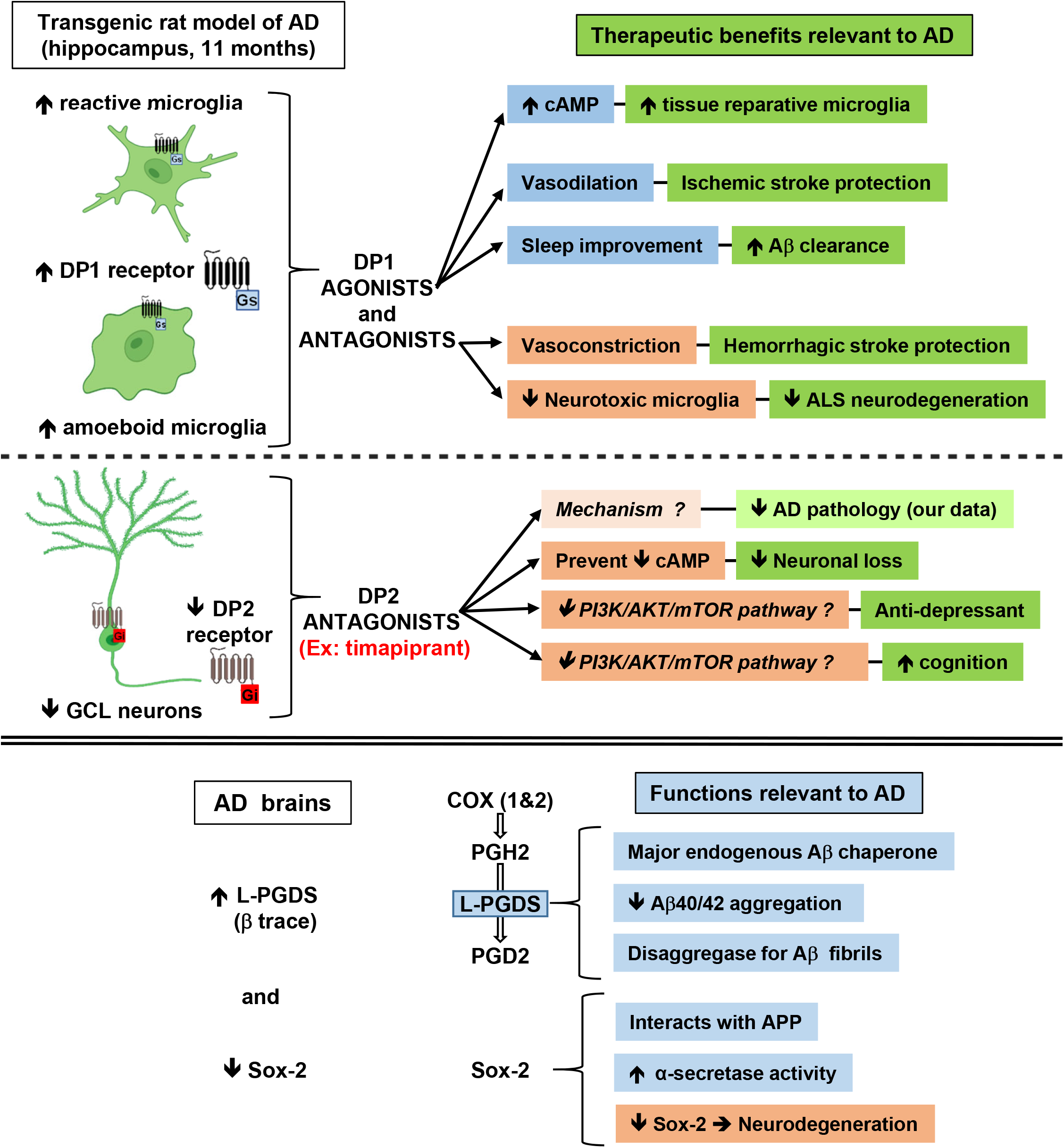
Scheme depicting the relevance of the PGD2 pathway to AD and its potential as a novel multitarget therapeutic for AD Tg-AD rats at 11 months of age exhibited a microglia shift in the hippocampus, from a neuroprotective state typical of ramified microglia to more of a neurotoxic and overactive state attributable to amoeboid microglia. DP1/microglia co-localization at the hippocampal hilar subregion followed the same pattern, increasing the most in amoeboid microglia. Tg-AD rats also displayed DP2 and neuronal losses in the hippocampus. In the brains of AD patients, L-PGDS, the major PGD2 synthase in the brain, is upregulated and Sox-2, a transcription factor, is downregulated. We propose that manipulating PGD2 signaling through, for example DP2 receptor antagonists such as timapiprant, could prevent/mitigate AD neuropathology. DP1 agonists via cAMP convert microglia to a tissue reparative phenotype, protect the brain from ischemic stroke by facilitating vasodilation, and promote sleep that drives Aβ clearance. DP1 antagonists protect against hemorrhagic stroke by limiting bleeding, and mitigate neurotoxic microglia levels. DP2 antagonists prevent neuronal loss, act as antidepressants and improve cognition, potentially via inhibition of the PI3K/AKT/mTOR pathway. L-PGDS, the major PGD2 synthase in the brain, also acts as an Aβ chaperone, inhibits Aβ40/42 aggregation, functions as a disaggregase by disassembling Aβ fibrils, and modulates the expression of Sox-2, a transcription factor. Sox-2 protects against AD neuropathology by interacting with APP and promoting α-secretase activity. Further details are presented in the discussion, based on our current results with the Tg-AD rat model and studies published by others. Figure partially created with BioRender.com.

In our studies with 11-month Tg-AD and WT rats, we confirmed that the DP2 receptor is highly expressed in hippocampal neurons, as previously shown by others [35]. DP2 is also expressed in astrocytes [43], but was not detected in microglia [13]. Since astrocyte levels were stable in the hippocampus of Tg-AD compared to WT rats, we focused our studies on DP2 and neuronal levels. Both neuronal and DP2 levels decreased significantly in a parallel manner in the GCL of the hippocampal DG region of Tg-AD rats compared with controls. The decline in DP2/neuronal levels could be tied to the rise in activated/amoeboid microglia. Thus, chronic PGD2 release as a result on enhanced neuroinflammation linked to AD, could on the one hand damage neurons via its DP2 receptor, and on the other hand increase the levels of activated/amoeboid microglia via its DP1 receptor. The changes in DP1 and DP2 levels that we report here are regional and specific, as they were only detected in the hilar subregion and GCL of the hippocampal DG (Fig. 10).

RNAseq analysis of 33 genes involved in the PGD2 and PGE2 pathways, demonstrated that mRNA transcript levels in whole (ventral and dorsal combined) hippocampal tissue were the highest for L-PGDS. Expression of L-PGDS is upregulated in AD phenotypes, correlates with Aβ plaque burden, and is associated with pathological traits of AD, but not with ALS or Parkinson’s disease [28, 30]. In our current studies, L-PGDS mRNA and protein levels were similar in WT and Tg-AD rats. However, whether changes occur in individual cell types and/or in specific hippocampal regions, like for the PGD2 receptors, remains to be determined.

L-PGDS also known as β-trace, is the primary PGD2 synthase in the brain, and is one of the most abundant (26 μg/ml, 3% of total) CSF proteins, second only to albumin [30, 64]. L-PGDS has a dual function, as it produces PGD2 and also acts as a lipophilic ligand-binding protein [64]. L-PGDS is a major endogenous Aβ chaperone that inhibits Aβ40/42 aggregation in vitro and in vivo, the latter when administered to mice intraventricularly infused with Aβ42 [28]. In vitro studies also demonstrated that L-PGDS acts as a disaggregase by disassembling Aβ fibrils [30]. In the PNS, L-PGDS contributes to myelination during development [63], and potentially acts as an antiinflammatory agent under conditions of peripheral nerve injury [22]. In the latter studies, L-PGDS modulated the expression of the transcription factor Sox-2, which in the CNS regulates oligodendrocyte proliferation and differentiation [71], and in the PNS is a negative regulator of myelination [20, 22]. Out of the 33 PG-associated genes that we focused on in our RNAseq analysis, Sox-2 expression was the third highest after L-PGDS and prostaglandin E synthase 3. Sox-2 is proposed to act as a protective factor in AD, as (1) it interacts with APP and mediates α-secretase activation in human cells, (2) its down-regulation in adult mouse brains induces neurodegeneration, and (3) its expression is downregulated in the brains of AD patients [55, 56]. Overall, more research is needed to establish whether modulating L-PDGS and Sox-2 has potential for preventing or treating AD.

Our results from treating Tg-AD rats with timapiprant reveal that manipulating PGD2 signaling with DP2 antagonists could potentially mitigate plaque load, neuronal loss and microglyosis, in addition to improving cognitive outcomes in AD patients (Fig. 10). In the brain, DP2 activation accelerates damage, as corroborated by studies with rat hippocampal neuronal and organotypic cultures in paradigms of glutamate toxicity [35, 36] or aluminum overload [39], and in a rat model of type 2 diabetes [70]. In the latter study, DP2 signaling promoted brain damage and inhibited autophagy by activating the PI3K/AKT/mTOR pathway [70]. Moreover, DP2 signaling mediates depression as well as cognitive dysfunction, supported by DP2-deficient mice exhibiting anti-depressant-like activity in a chronic corticosterone-induced model of depression [47], and improved cognition in an NMDA receptor antagonist-induced model of cognitive dysfunction [48]. These findings support that DP2 signaling has a negative impact on emotion and cognition. Thus, selective DP2 receptor antagonists may represent an encouraging option for treating some types of brain disorders. However, further studies are necessary to establish the long-term safety and benefits of these drugs that could be used as a monotherapy or in combination with other therapies aimed, for example, at reducing amyloid plaque burden in AD.

## 5. Conclusions

In conclusion, at the periphery, PGD2 is an established inflammatory mediator [42], and its effects include enhancing vascular permeability [21], modulating chemotaxis [26] and antigen presentation [25], as well as vasodilatation, bronchoconstriction, inhibition of platelet aggregation, glycogenolysis, allergic reaction mediation, and intraocular pressure reduction (reviewed in [5]), as well as potentially resolving peripheral nerve injury [22]. In the CNS, PGD2 regulates sleep induction, body temperature, olfactory function, nociception, neuromodulation, and protects the brain from ischemic stroke (reviewed in [5]). Our current data suggest that, as an alternative to NSAIDs and as a novel approach for treating neuroinflammation, manipulating PGD2 signaling with for example DP2 receptor antagonists, could have a significant translational and multifactorial potential as a therapeutic for AD.

## Abbreviations

Aβ: amyloid β
AD: Alzheimer’s disease
CA: cornu ammonis
COX: cyclooxygenase
DG: dentate gyrus
DP1: prostaglandin D2 receptor 1
DP2: prostaglandin D2 receptor 2
DG: dentate gyrus
FDR: false discovery rate
FL-APP: full length amyloid precursor protein
GAPDH: glyceraldehyde 3-phosphate dehydrogenase
GFAP: Glial fibrillary acidic protein
GCL: granular cell layer
HL: hilar
Iba1: ionized calcium binding adaptor molecule 1
L-PGDS: lipocalin-type prostaglandin D synthase
NeuN: neuronal nuclei, neuronal marker
PG: prostaglandin
PPARγ: peroxisome proliferator activated receptor gamma
SB: subiculum
SEM: standard error of the mean
RPM: reads per million
Sox-2: SRY-box transcription factor 2
Tg-AD: transgenic rat model of Alzheimer’s disease
WT: wild type

## 6. Acknowledgments

This work was supported in part by NIH/NIA R01AG057555 to LX, NIH training grants 5T32HL135465-A1 to support CHW and R25GM060665 to support GO, and the City University of New York (Ph.D. program in Biochemistry, Graduate Center). We thank the technical support of Osama Chaudry^a^, Rushna Snetha^a^, Aminoor Rashid^a^ (all Master’s students), and Amber Alliger^b^ (Lecturer) in the Departments of Biological Sciences^a^ and Psychology^b^ at Hunter College, CUNY. We also thank Ms. Lisa Bleyle and Dr. Dennis Koop for the assistance with the prostaglandin analysis conducted in the Bioanalytical Shared Resource/Pharmacokinetics Core. The facility is part of the University Shared Resource Program at Oregon Health

## 7. CRediT author statement

**Charles H. Wallace,** Investigation, Formal analysis, Writing-Original draft preparation, Visualization. **Giovanni Oliveros**: Investigation. **Peter A Serrano, Patricia Rockwell:** Validation, Writing-Reviewing and Editing. **Lei Xie:** Reviewing, Funding acquisition. **Maria E. Figueiredo-Pereira:** Conceptualization, Writing-Original draft preparation.

## 8. Supplemental material

### 8.1. Tables

Table 1: DP1 levels (% signal) across hippocampal regions

Table 2: Microglia levels (Iba1+, counts/nm^2^) across hippocampal regions

Table 3: DP1 and microglia (Iba1+) co-localization across hippocampal regions

Table 4: Astrocyte levels (GFAP, % signal) across hippocampal regions

Table 5: Neuronal levels (NeuN, % signal) across hippocampal regions

Table 6: DP2 levels (% signal) across hippocampal regions

Table 7: DP2 and neuronal (NeuN+) co-localization across hippocampal regions

Table 8: RNA sequence analysis for selected genes of the prostaglandin pathway in hippocampal tissue

Table 9: Semi-quantification of protein expression in hippocampal tissue

Table 10: Antibodies used for IHC and WB analyses of hippocampal tissue

### 8.2. Figures

Fig. 1: Subregions within the dentate gyrus

Fig. 2: RNAseq – PG biosynthesis and metabolism

Fig. 3: RNAseq – PGD2 and PGE2 synthases

Fig. 4: RNAseq – PGD2 and PGJ2 receptors

Fig. 5: RNAseq – PPARγ receptor activators and Sox-2

Fig. 6: RNAseq – PGE2 receptors

Fig. 7: Western blot analyses

Fig. 8: Full blots – DP1, DP2, L-PGDS

Fig. 9: Full blots – FL-APP and Aβ

Fig. 10: IHC hippocampal images for manuscript Fig. 9

