## Supplemental Tables & Figures for "Prostaglandin D2 pathway in a transgenic rat model of Alzheimer’s disease: therapeutic potential of timapiprant a DP2 antagonist"

**Supplemental Table 1** DP1 levels (% signal) across hippocampal regions

| | WT mean ± SEM n = 12 | Tg-AD mean ± SEM n = 11 | *p*-value | *t*-statistics |
|---|---|---|---|---|
| **SB** | 3.88 ± 0.16 | 4.05 ± 0.23 | 0.27 | t=0.15, df=17.08 |
| **CA1** | 3.50 ± 0.20 | 3.97 ± 0.22 | 0.06 | t=1.58, df=20.76 |
| **CA3** | 3.91 ± 0.16 | 4.24 ± 0.25 | 0.13 | t=1.14, df=17.23 |
| **DG** | 4.33 ± 0.23 | 4.63 ± 0.16 | 0.15 | t=1.06, df=19.31 |
| **HL** | 5.22 ± 0.22 | 5.34 ± 0.22 | 0.36 | t=0.36, df=20.99 |

Values represent the percent of the signal (DP1+) detected within a specific area (= 100%, $\mu m^2$) as explained under materials and methods. Abbreviations: DP1, prostaglandin D2 receptor 1; WT, wild type; Tg-AD, transgenic rat model of Alzheimer's disease; SEM, standard error of the mean; SB, subiculum; CA, cornu ammonis; DG, dentate gyrus; HL, hilar.

**Supplemental Table 2** Microglia levels (Iba1+, counts/$nm^2$) across hippocampal regions

| | WT mean ± SEM n = 12 | Tg-AD mean ± SEM n = 11 | *p*-value | *t*-statistics |
|---|---|---|---|---|
| **SB** | 214.90 ± 7.75 | 235.80 ± 9.08 | 0.05 | t=1.75, df=20.16 |
| **CA1** | 231.10 ± 8.65 | 263.00 ± 11.37 | 0.02 | t=2.23, df=19.10 |
| **CA3** | 246.10 ± 7.60 | 269.20 ± 12.27 | 0.06 | t=1.60, df=16.89 |
| **DG** | 226.10 ± 6.86 | 268.20 ± 11.20 | 0.003 | t=3.20, df=16.76 |
| **HL** | 40.70 ± 2.47 | 58.95 ± 5.24 | 0.003 | t=3.15, df=14.28 |

Values represent the microglia counts (Iba1+) detected within a specific area ($nm^2$) as explained under materials and methods. Abbreviations: Iba1, ionized calcium binding adaptor molecule 1; WT, wild type; Tg-AD, transgenic rat model of Alzheimer's disease; SEM, standard error of the mean; SB, subiculum; CA, cornu ammonis; DG, dentate gyrus; HL, hilar.

**Supplemental Table 3** DP1 and microglia (Iba1+) co-localization across hippocampal regions

| | WT mean ± SEM n = 12 | Tg-AD mean ± SEM n = 11 | *p*-value | *t*-statistics |
|---|---|---|---|---|
| **SB** | 171.70 ± 11.07 | 178.30 ± 7.08 | 0.31 | t=0.50, df=18.44 |
| **CA1** | 178.60 ± 11.93 | 189.30 ± 9.21 | 0.24 | t=0.71, df=20.15 |
| **CA3** | 206.50 ± 9.71 | 218.50 ± 10.00 | 0.20 | t=0.86, df=20.87 |
| **DG** | 190.50 ± 8.70 | 226.40 ± 12.61 | 0.02 | t=2.35, df=18.07 |
| **HL** | 35.34 ± 2.26 | 52.54 ± 5.27 | 0.005 | t=2.00, df=13.60 |

Values represent microglia counts (Iba1+) co-localized with DP1 signal within a specific area ($nm^2$) as explained under materials and methods. Abbreviations: DP1, prostaglandin D2 receptor 1; Iba1, ionized calcium binding adaptor molecule 1; WT, wild type; Tg-AD, transgenic rat model of Alzheimer's disease; SEM, standard error of the mean; SB, subiculum; CA, cornu ammonis; DG, dentate gyrus; HL, hilar.

**Supplemental Table 4** Astrocyte levels (GFAP, % signal) across hippocampal regions

| | WT mean ± SEM n = 12 | Tg-AD mean ± SEM n = 11 | *p*-value | *t*-statistics |
|---|---|---|---|---|
| **SB** | 4.42 ± 0.13 | 4.56 ± 0.20 | 0.28 | t=0.58, df=17.13 |
| **CA1** | 4.36 ± 0.11 | 4.61 ± 0.17 | 0.11 | t=1.26, df=17.37 |
| **CA3** | 4.45 ± 0.15 | 4.65 ± 0.16 | 0.23 | t=0.75, df=20.47 |
| **DG** | 4.47 ± 0.12 | 4.71 ± 0.12 | 0.08 | t=1.44, df=20.81 |
| **HL** | 4.65 ± 0.21 | 4.99 ± 0.20 | 0.13 | t=1.15, df=21.00 |

Values represent the percent of the signal (GFAP+) detected within a specific area (= 100%, $\mu m^2$) as explained under materials and methods. Abbreviations: GFAP, Glial fibrillary acidic protein; WT, wild type; Tg-AD, transgenic rat model of Alzheimer's disease; SEM, standard error of the mean; SB, subiculum; CA, cornu ammonis; DG, dentate gyrus; HL, hilar.

**Supplemental Table 5** Neuronal levels (NeuN, % signal) across hippocampal regions

| | WT mean ± SEM n = 12 | Tg-AD mean ± SEM n = 11 | *p*-value | *t*-statistics |
|---|---|---|---|---|
| **SB** | 5.55 ± 0.30 | 5.55 ± 0.41 | 0.50 | t=0.004, df=18.67 |
| **CA1** | 4.13 ± 0.13 | 4.09 ± 0.21 | 0.44 | t=0.16, df=17.21 |
| **CA3** | 6.38 ± 0.19 | 6.11± 0.22 | 0.18 | t=0.95, df=20.10 |
| **DG** | 6.29 ± 0.37 | 5.32 ± 0.38 | 0.04 | t=1.83, df=20.82 |
| **HL** | 6.70 ± 0.25 | 5.76 ± 0.49 | 0.05 | t=1.71, df=14.96 |

Values represent the percent of the signal (NeuN+) detected within a specific area (= 100%, $\mu m^2$) as explained under materials and methods. Abbreviations: NeuN, neuronal nuclei, neuronal marker; WT, wild type; Tg-AD, transgenic rat model of Alzheimer's disease; SEM, standard error of the mean; SB, subiculum; CA, cornu ammonis; DG, dentate gyrus; HL, hilar.

**Supplemental Table 6** DP2 levels (% signal) across hippocampal regions

| | WT mean ± SEM n = 12 | Tg-AD mean ± SEM n = 11 | *p*-value | *t*-statistics |
|---|---|---|---|---|
| **SB** | 4.35 ± 0.20 | 4.36 ± 0.21 | 0.29 | t=0.05, df=20.75 |
| **CA1** | 3.48 ± 0.16 | 4.47 ± 0.25 | 0.002 | t=3.36, df=17.62 |
| **CA3** | 3.85 ± 0.18 | 4.21 ± 0.14 | 0.07 | t=1.56, df=20.30 |
| **DG** | 4.62 ± 0.17 | 4.87 ± 0.13 | 0.13 | t=1.14, df=20.30 |
| **HL** | 5.29 ± 0.14 | 5.44 ± 0.13 | 0.21 | t=0.83, df=20.97 |

Values represent the percent of the signal (DP2+) detected within a specific area (= 100%, $\mu m^2$) as explained under materials and methods. Abbreviations: DP2, prostaglandin D2 receptor 2; WT, wild type; Tg-AD, transgenic rat model of Alzheimer's disease; SEM, standard error of the mean; SB, subiculum; CA, cornu ammonis; DG, dentate gyrus; HL, hilar.

**Supplemental Table 7** DP2 and neuronal (NeuN+) co-localization across hippocampal regions

| | **WT mean ± SEM n = 12** | **Tg-AD mean ± SEM n = 11** | ***p*-value** | ***t*-statistics** |
|---|---|---|---|---|
| **SB** | 51.69± 5.17 | 47.96± 4.29 | 0.29 | t=0.56, df=20.61 |
| **CA1** | 50.26 ± 4.11 | 48.21 ± 3.71 | 0.36 | t=0.37, df=20.94 |
| **CA3** | 47.33± 3.29 | 44.44± 3.85 | 0.29 | t=0.57, df=20.17 |
| **DG** | 46.75± 3.13 | 47.14± 3.89 | 0.47 | t=0.08, df=19.65 |
| **HL** | 67.27 ± 2.85 | 67.48 ± 4.29 | 0.48 | t=0.04, df=17.66 |

Values represent % co-localized NeuN and DP2 signals within a specific area ($nm^2$) as explained under materials and methods. Abbreviations: DP2, prostaglandin D2 receptor 2; NeuN, neuronal nuclei, neuronal marker; WT, wild type; Tg-AD, transgenic rat model of Alzheimer's disease; SEM, standard error of the mean; SB, subiculum; CA, cornu ammonis; DG, dentate gyrus; HL, hilar.

**Supplemental Table 8** RNA sequence analysis for selected prostaglandin pathway genes

| a MALES | | | | | | |
|---|---|---|---|---|---|---|
| **Gene symbol** | **Function/Name** | **Fold change** | ***p* Value** | **FDR** | **Tg-AD mean ± SEM** | **WT mean ± SEM** |
| | **PROSTAGLANDIN D2 and J2** | | | | | |
| **Ptgds** | prostaglandin D2 synthase (brain) | 1.83 | 0.212 | 1 | 358.94 ± 53.18 | 258.16 ± 23.38 |
| **Sox2** | SRY (sex determining region Y)-box 2 | -1.29 | 0.011 | 0.550 | 88.47 ± 8.08 | 113.72 ± 5.36 |
| **Ppard** | peroxisome proliferator-activated receptor delta | -1.18 | 0.110 | 1 | 42.23 ± 3.74 | 49.71 ± 3.26 |
| **Pprc1** | peroxisome proliferator-activated receptor gamma, coactivator-related 1 | 1.06 | 0.474 | 1 | 28.32 ± 0.65 | 26.71 ± 0.77 |
| **Ppargc1a** | peroxisome proliferator-activated receptor gamma, coactivator 1 alpha | -1.08 | 0.507 | 1 | 15.53 ± 0.55 | 16.76 ± 0.81 |
| **Ppargc1b** | peroxisome proliferator-activated receptor gamma, coactivator 1 beta | 1.00 | 1.000 | 1 | 4.93 ± 0.26 | 4.93 ± 0.26 |
| **Ppara** | peroxisome proliferator activated receptor alpha | -1.09 | 0.600 | 1 | 3.7 ± 0.37 | 4.04 ± 0.25 |
| **Hpgds** | hematopoietic prostaglandin D synthase | -1.04 | 0.890 | 1 | 1.19 ± 0.32 | 1.24 ± 0.15 |
| **Ptgdrl** | prostaglandin D2 receptor-like (in rat, orthologous to human DP1) | -1.87 | 0.296 | 1 | 0.34 ± 0.10 | 0.64 ± 0.13 |
| **Ptgdr2** | prostaglandin D2 receptor 2 (DP2) | -1.16 | 0.667 | 1 | 0.51 ± 0.18 | 0.59 ± 0.06 |
| **Pparg** | peroxisome proliferator-activated receptor gamma | 1.07 | 0.872 | 1 | 0.26 ± 0.03 | 0.24 ± 0.02 |
| **Ptgdr** | prostaglandin D2 receptor (DP1) | 3.30 | 0.164 | 1 | 0.11 ± 0.05 | 0.03 ± 0.03 |
| | **PROSTAGLANDIN E2** | | | | | |
| **Ptges3** | prostaglandin E synthase 3 (cytosolic) | -1.03 | 0.748 | 1 | 156.20 ± 8.97 | 160.45 ± 5.78 |
| **Ptges2** | prostaglandin E synthase 2 | -1.00 | 0.975 | 1 | 46.14 ± 0.98 | 46.22 ± 0.93 |
| **Ptges3l1** | prostaglandin E synthase 3-like 1 | -1.06 | 0.582 | 1 | 16.29 ± 0.87 | 17.32 ± 1.22 |
| **Ptger1** | prostaglandin E receptor 1 (subtype EP1) | -1.31 | 0.027 | 0.825 | 6.59 ± 0.17 | 8.65 ± 0.54 |
| **Ptger3** | prostaglandin E receptor 3 (subtype EP3) | 1.56 | 0.095 | 1 | 1.58 ± 0.20 | 1.01 ± 0.18 |
| **Ptger2** | prostaglandin E receptor 2 (subtype EP2) | -1.02 | 0.946 | 1 | 0.97 ± 0.05 | 0.99 ± 0.21 |
| **Ptges** | prostaglandin E synthase | 1.15 | 0.712 | 1 | 1.06 ± 0.29 | 0.93 ± 0.26 |
| **Ptger4** | prostaglandin E receptor 4 (subtype EP4) | 1.94 | 0.082 | 1 | 0.44 ± 0.12 | 0.23 ± 0.06 |
| | **THROMBOXANE** | | | | | |
| **Tbxas1** | thromboxane A synthase 1, platelet | -1.02 | 0.906 | 1 | 4.51 ± 0.81 | 4.60 ± 0.36 |
| **Tbxa2r** | thromboxane A2 receptor | -1.51 | 0.076 | 1 | 1.01 ± 0.14 | 1.52 ± 0.16 |
| | **PHOSPHOLIPASES, CYCLOOXYGENASES & RELATED** | | | | | |
| **Ptgr2** | prostaglandin reductase 2 | -1.16 | 0.091 | 1 | 88.12 ± 5.78 | 102.54 ± 6.78 |
| **Ptgs2** | prostaglandin-endoperoxide synthase 2 (COX-2) | -1.20 | 0.473 | 1 | 18.51 ± 2.86 | 22.28 ± 5.27 |
| **Ptgs1** | prostaglandin-endoperoxide synthase 1 (COX-1) | -1.13 | 0.328 | 1 | 17.52 ± 2.15 | 19.86 ± 2.13 |
| **Pla2g4a** | phospholipase A2, group IVA (cytosolic, calcium-dependent) | -1.02 | 0.877 | 1 | 7.70 ± 0.43 | 7.87 ± 0.65 |
| **Hpgd** | hydroxyprostaglandin dehydrogenase 15 (NAD) | 1.17 | 0.562 | 1 | 2.97 ± 0.57 | 2.53 ± 0.45 |
| **Ptgr1** | prostaglandin reductase 1 | 1.03 | 0.939 | 1 | 1.07 ± 0.14 | 1.05 ± 0.23 |
| | **OTHERS** | | | | | |
| **Ptgfrn** | prostaglandin F2 receptor inhibitor | -1.06 | 0.702 | 1 | 40.05 ± 5.40 | 42.33 ± 5.36 |
| **Ptgis** | prostaglandin I2 (prostacyclin) synthase | 1.26 | 0.508 | 1 | 1.76 ± 0.29 | 1.40 ± 0.21 |
| **Ptgfr** | prostaglandin F receptor | 1.90 | 0.246 | 1 | 1.22 ± 0.67 | 0.65 ± 0.21 |
| **Ptgir** | prostaglandin I2 (prostacyclin) receptor (IP) | -1.65 | 0.313 | 1 | 0.11 ± 0.05 | 0.19 ± 0.04 |
| **Cbr1** | carbonyl reductase 1 | 1.10 | 0.250 | 1 | 38.78 ± 1.20 | 35.15 ± 2.47 |

| b FEMALES | | | | | | |
|---|---|---|---|---|---|---|
| **Gene symbol** | **Function/Name** | **Fold change** | **P Value** | **FDR** | **Tg-AD mean ± SEM** | **WT mean ± SEM** |
| | **PROSTAGLANDIN D2 and J2** | | | | | |
| **Ptgds** | prostaglandin D2 synthase (brain) | -1.49 | 0.070 | 1 | 189.32 ± 8.05 | 282.71 ± 94.63 |
| **Sox2** | SRY (sex determining region Y)-box 2 | -1.06 | 0.636 | 1 | 75.22 ± 5.32 | 79.43 ± 8.82 |
| **Ppard** | peroxisome proliferator-activated receptor delta | -1.01 | 0.904 | 1 | 37.98 ± 0.93 | 38.41 ± 0.50 |
| **Pprc1** | peroxisome proliferator-activated receptor gamma, coactivator-related 1 | 1.07 | 0.471 | 1 | 32.48 ± 1.26 | 30.34 ± 0.93 |
| **Ppargc1a** | peroxisome proliferator-activated receptor gamma, coactivator 1 alpha | -1.06 | 0.615 | 1 | 19.07 ± 1.01 | 20.22 ± 1.51 |
| **Ppargc1b** | peroxisome proliferator-activated receptor gamma, coactivator 1 beta | 1.01 | 0.950 | 1 | 6.67 ± 0.89 | 6.58 ± 0.51 |
| **Ppara** | peroxisome proliferator activated receptor alpha | -1.01 | 0.956 | 1 | 3.63 ± 0.16 | 3.66 ± 0.39 |
| **Hpgds** | hematopoietic prostaglandin D synthase | 1.31 | 0.408 | 1 | 1.48 ± 0.30 | 1.13 ± 0.19 |
| **Ptgdrl** | prostaglandin D2 receptor-like (in rat, orthologous to human DP1) | 1.65 | 0.304 | 1 | 0.71 ± 0.11 | 0.43 ± 0.18 |
| **Ptgdr2** | prostaglandin D2 receptor 2 (DP2) | -1.17 | 0.676 | 1 | 0.35 ± 0.08 | 0.41 ± 0.03 |
| **Pparg** | peroxisome proliferator-activated receptor gamma | 1.13 | 0.769 | 1 | 0.28 ± 0.03 | 0.24 ± 0.06 |
| **Ptgdr** | prostaglandin D2 receptor (DP1) | -3.01 | 0.187 | 1 | 0.03 ± 0.03 | 0.09 ± 0.06 |
| | **PROSTAGLANDIN E2** | | | | | |
| **Ptges3** | prostaglandin E synthase 3 (cytosolic) | 1.02 | 0.779 | 1 | 169.56 ± 3.85 | 165.83 ± 3.29 |
| **Ptges2** | prostaglandin E synthase 2 | 1.03 | 0.760 | 1 | 47.98 ± 2.39 | 46.52 ± 2.50 |
| **Ptges3l1** | prostaglandin E synthase 3-like 1 | -1.01 | 0.926 | 1 | 16.27 ± 0.70 | 16.46 ± 0.52 |
| **Ptger1** | prostaglandin E receptor 1 (subtype EP1) | -1.02 | 0.902 | 1 | 5.38 ± 0.36 | 5.51 ± 0.31 |
| **Ptger3** | prostaglandin E receptor 3 (subtype EP3) | -1.09 | 0.739 | 1 | 1.41 ± 0.09 | 1.54 ± 0.15 |
| **Ptger2** | prostaglandin E receptor 2 (subtype EP2) | 1.00 | 0.994 | 1 | 1.14 ± 0.29 | 1.14 ± 0.04 |
| **Ptges** | prostaglandin E synthase | -1.86 | 0.089 | 1 | 0.62 ± 0.16 | 1.18 ± 0.21 |
| **Ptger4** | prostaglandin E receptor 4 (subtype EP4) | -1.11 | 0.780 | 1 | 0.35 ± 0.07 | 0.39 ± 0.04 |
| | **THROMBOXANE** | | | | | |
| **Tbxas1** | thromboxane A synthase 1, platelet | 1.19 | 0.382 | 1 | 3.83 ± 0.14 | 3.21 ± 0.21 |
| **Tbxa2r** | thromboxane A2 receptor | -1.23 | 0.549 | 1 | 1.00 ± 0.22 | 1.24 ± 0.34 |
| | **PHOSPHOLIPASES, CYCLOOXYGENASES & RELATED** | | | | | |
| **Ptgr2** | prostaglandin reductase 2 | 1.03 | 0.757 | 1 | 72.66 ± 3.39 | 70.62 ± 3.63 |
| **Ptgs2** | prostaglandin-endoperoxide synthase 2 (COX-2) | -1.09 | 0.604 | 1 | 21.64 ± 2.25 | 23.62 ± 3.51 |
| **Ptgs1** | prostaglandin-endoperoxide synthase 1 (COX-1) | 1.07 | 0.612 | 1 | 17.71 ± 0.71 | 16.61 ± 1.75 |
| **Pla2g4a** | phospholipase A2, group IVA (cytosolic, calcium-dependent) | -1.09 | 0.599 | 1 | 6.73 ± 0.43 | 7.36 ± 0.62 |
| **Hpgd** | hydroxyprostaglandin dehydrogenase 15 (NAD) | 1.26 | 0.358 | 1 | 3.64 ± 0.13 | 2.87 ± 0.57 |
| **Ptgr1** | prostaglandin reductase 1 | 1.35 | 0.361 | 1 | 0.91 ± 0.12 | 0.68 ± 0.12 |
| | **OTHERS** | | | | | |
| **Ptgfrn** | prostaglandin F2 receptor inhibitor | -1.42 | 0.027 | 1 | 32.58 ± 0.97 | 46.23 ± 9.27 |
| **Ptgis** | prostaglandin I2 (prostacyclin) synthase | -1.13 | 0.700 | 1 | 1.12 ± 0.24 | 1.27 ± 0.17 |
| **Ptgfr** | prostaglandin F receptor | -1.08 | 0.832 | 1 | 0.51 ± 0.09 | 0.55 ± 0.11 |
| **Ptgir** | prostaglandin I2 (prostacyclin) receptor (IP) | -1.19 | 0.734 | 1 | 0.18 ± 0.03 | 0.21 ± 0.09 |
| **Cbr1** | carbonyl reductase 1 | -1.11 | 0.377 | 1 | 38.02 ± 0.70 | 42.14 ± 4.51 |

Within each functional group, genes are listed in decreasing order of expression levels (RPM mean ± SEM) for Tg-AD (n = 5) and WT (n = 5) male (A) and female (B) rats. Gene functions/names, fold change, *P* values, and FDR are also included. Abbreviations: WT, wild type; Tg-AD, transgenic rat model of Alzheimer's disease; SEM, standard error of the mean; RPM, reads per million; FDR, false discovery rates.

**Supplemental Table 9** Semi-quantification of protein expression in hippocampal tissue

| | **WT mean ± SEM n = 6** | **Tg-AD mean ± SEM n = 6** | ***p*-value** | ***t*-statistics** |
|---|---|---|---|---|
| **FL-APP/actin** | 1.00 ± 0.22 | 5.62 ± 0.60 | <0.001 | t=7.23, df=6.30 |
| **Aβ/GAPDH** | 1.00 ± 0.20 | 32.60 ± 8.73 | 0.008 | t=3.62, df=5.01 |
| **COX-2/tubulin** | 1.00 ± 0.18 | 1.68 ± 0.38 | 0.07 | t=1.63, df=7.15 |
| **DP1/GAPDH** | 1.00 ± 0.08 | 1.36 ± 0.08 | 0.16 | t=1.04, df=8.96 |
| **DP2/GAPDH** | 1.00 ± 0.16 | 0.79 ± 0.09 | 0.15 | t=1.13, df=7.95 |
| **L-PGDS/GAPDH** | 1.00 ± 0.08 | 1.09 ± 0.14 | 0.30 | t=0.55, df=8.05 |
| **PPARγ/actin** | 1.00 ± 0.18 | 0.68 ± 0.14 | 0.10 | t=1.40, df=9.53 |
| **Sox-2/actin** | 1.00 ± 0.32 | 0.72 ± 0.27 | 0.24 | t=0.73, df=9.19 |

Values represent the respective protein amounts detected in hippocampal homogenates. Abbreviations: WT, wild type; Tg-AD, transgenic rat model of Alzheimer's disease; SEM, standard error of the mean; FL-APP, full length amyloid precursor protein; Aβ, amyloid β; GAPDH, glyceraldehyde 3-phosphate dehydrogenase; COX-2, cyclooxygenase-2; DP1, prostaglandin D2 receptor 1; DP2, prostaglandin D2 receptor 2; L-PGDS, lipocalin-type prostaglandin D synthase; PPARγ, peroxisome proliferator activated receptor gamma; Sox-2, SRY-box transcription factor 2.

**Supplemental Table 10** Antibodies used for IHC and WB analyses of hippocampal tissue

| Antibody | Company | Catalog Number | Species/Type | Dilution | Assay |
|---|---|---|---|---|---|
| ***PRIMARIES*** | | | | | |
| **Aβ (4G8)** | Biolegend | #800708 | Mouse Monoclonal | 1:1000 | IHC |
| **Aβ (6E10)** | Biolegend | #SIG-39320 | Mouse Monoclonal | 1:2000 | WB |
| **COX-2 (D5H5)** | Cell Signaling | #12282 | Rabbit Monoclonal | 1:1000 | WB |
| **DP1** | Cayman Chemical | #101640 | Rabbit Polyclonal | 1:200 for both | IHC, WB |
| **DP2** | Invitrogen | #PA5-20332 | Rabbit Polyclonal | 1:1000 for both | IHC, WB |
| **FL-APP (22C11)** | Millipore Sigma | #MAB348 | Mouse Monoclonal | 1:2000 | WB |
| **GAPDH (6C5)** | Millipore Sigma | #MAB374 | Mouse Monoclonal | 1:2000 | WB |
| **GFAP (GA5)** | Millipore Sigma | #MAB360 | Mouse Monoclonal | 1:1000 | IHC |
| **Iba1** | Synaptic Systems | #234006 | Chicken Polyclonal | 1:500 | IHC |
| **L-PGDS** | Abcam | #ab182141 | Rabbit Polyclonal | 1:1000 | WB |
| **NeuN** | Millipore Sigma | #ABN91 | Chicken Polyclonal | 1:500 | IHC |
| **PPARγ** | Abcam | #ab209350 | Rabbit Polyclonal | 1:1000 | WB |
| **Sox-2** | Abcam | #ab97959 | Rabbit Polyclonal | 1:2000 | WB |
| **β-Actin (AC-74)** | Sigma-Aldrich | #A2228 | Mouse Monoclonal | 1:10000 | WB |
| **β-Tubulin (TUBB3)** | Covance | #MMS-435P | Mouse Monoclonal | 1:10000 | WB |
| ***SECONDARIES*** | | | | | |
| **Alexa Fluor 488, Donkey anti-Mouse IgG (H+L)** | ThermoFisher | #A-21202 | Mouse Secondary | 1:250 | IHC |
| **Alexa Fluor 488, Goat anti-Chicken IgY (H+L)** | ThermoFisher | #A-11039 | Chicken Secondary | 1:500 | IHC |
| **Alexa Fluor 568, Goat anti-Rabbit IgG (H+L)** | ThermoFisher | #A-11011 | Rabbit Secondary | 1:250 | IHC |
| **Goat Anti-Rabbit IgG (H + L)-HRP** | Cell Signaling | #7074 | Rabbit Secondary | 1:3000-10000 | WB |
| **Hoarse Anti-Mouse IgG (H + L)-HRP** | Cell Signaling | #7076 | Mouse Secondary | 1:3000-10000 | WB |

Abbreviations: IHC, immunohistochemistry; WB, western blot; Aβ, amyloid β; COX-2, cyclooxygenase-2; DP1, prostaglandin D2 receptor 1; DP2, prostaglandin D2 receptor 2; FL-APP, full length amyloid precursor protein; GAPDH, glyceraldehyde 3-phosphate dehydrogenase; GFAP, glial fibrillary acidic protein; Iba1, ionized calcium binding adaptor molecule 1; NeuN, neuronal nuclei; neuronal marker; PPARγ, peroxisome proliferator activated receptor gamma; Sox-2, SRY-box transcription factor 2.

**Supplemental Fig. 1**

Subregions within the dentate gyrus (NeuN IHC for a WT female rat)

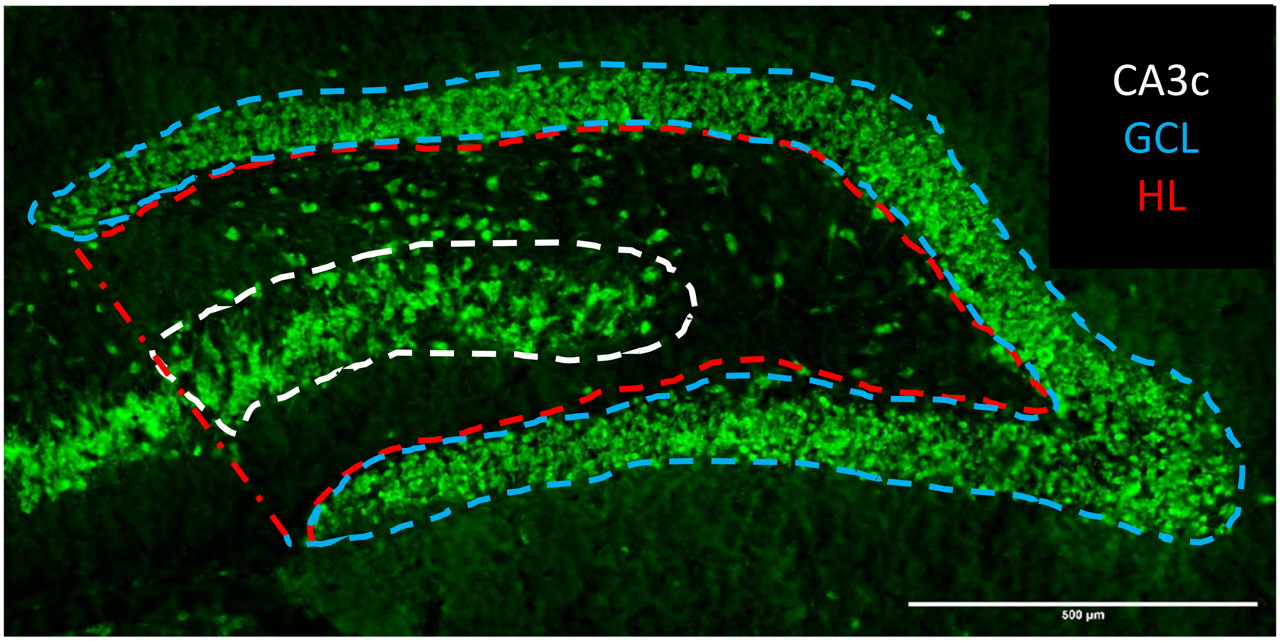


HL, hilar; GCL, granular cell layer, CA, cornu ammonis

### Supplemental Fig. 2

RNAseq – PG biosynthesis and metabolism

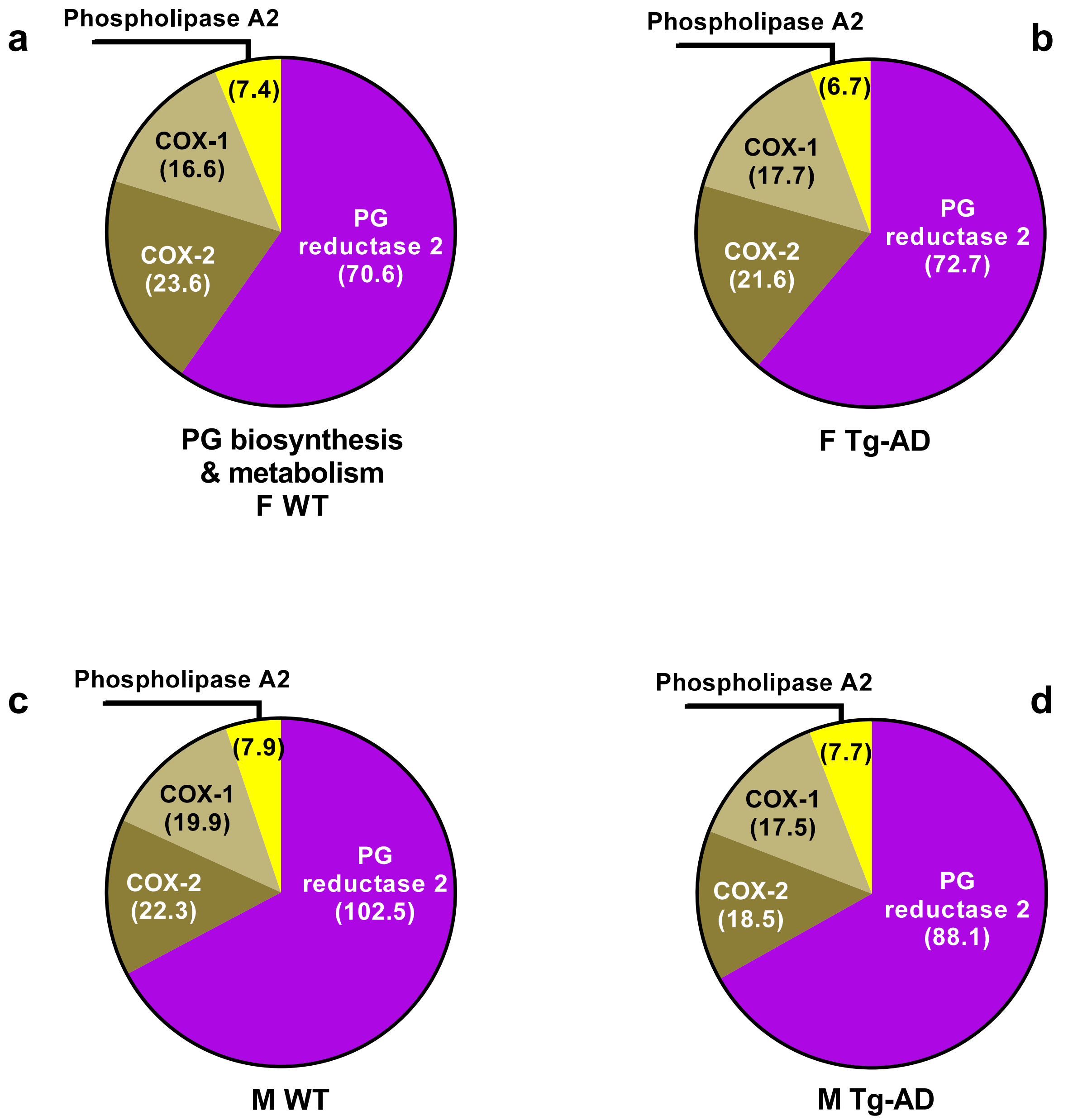

### Supplemental Fig. 3

RNAseq – PGD2 and PGE2 synthases

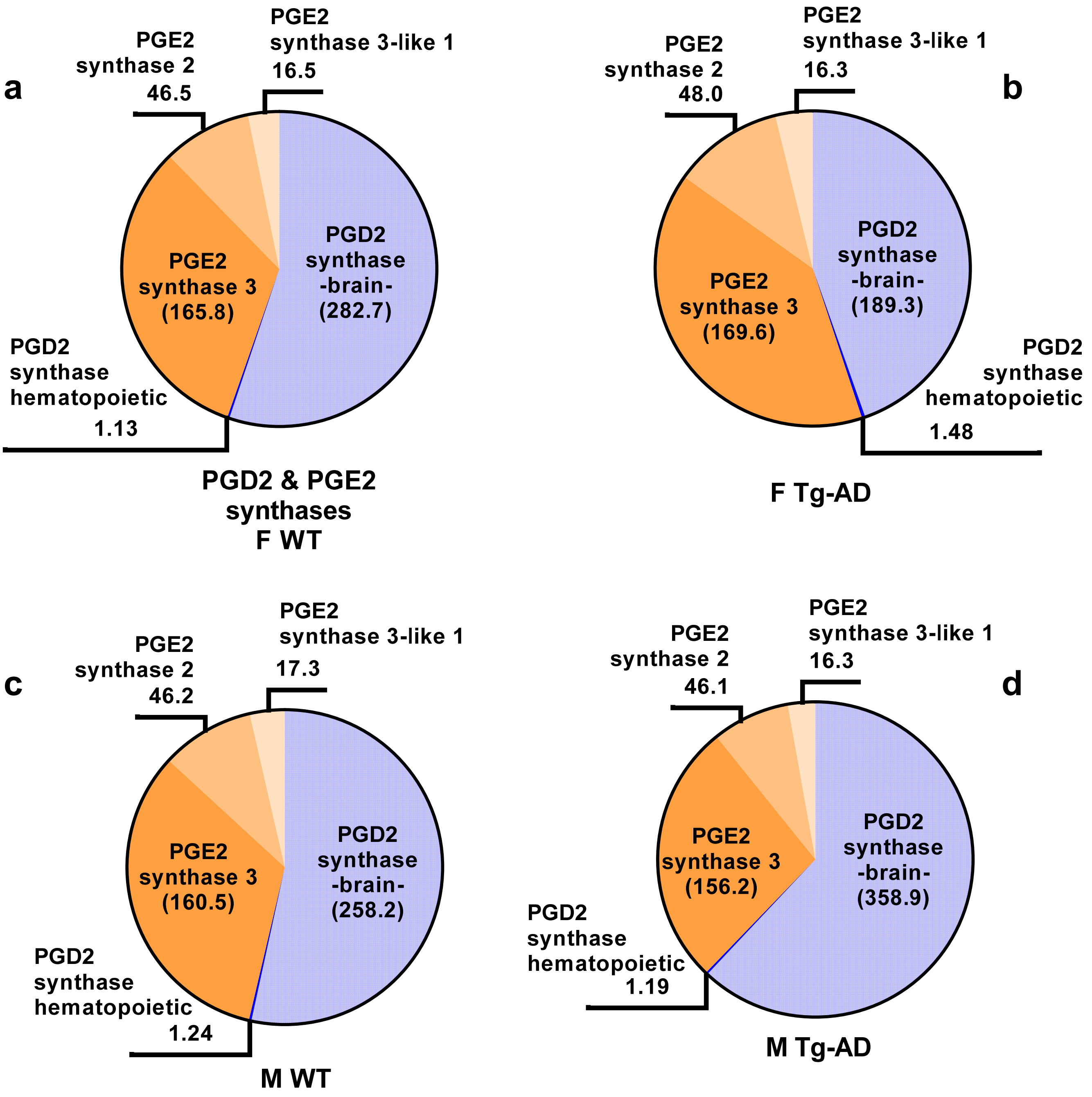

**Supplemental Fig. 4**

RNAseq – PGD2 and PGJ2 receptors

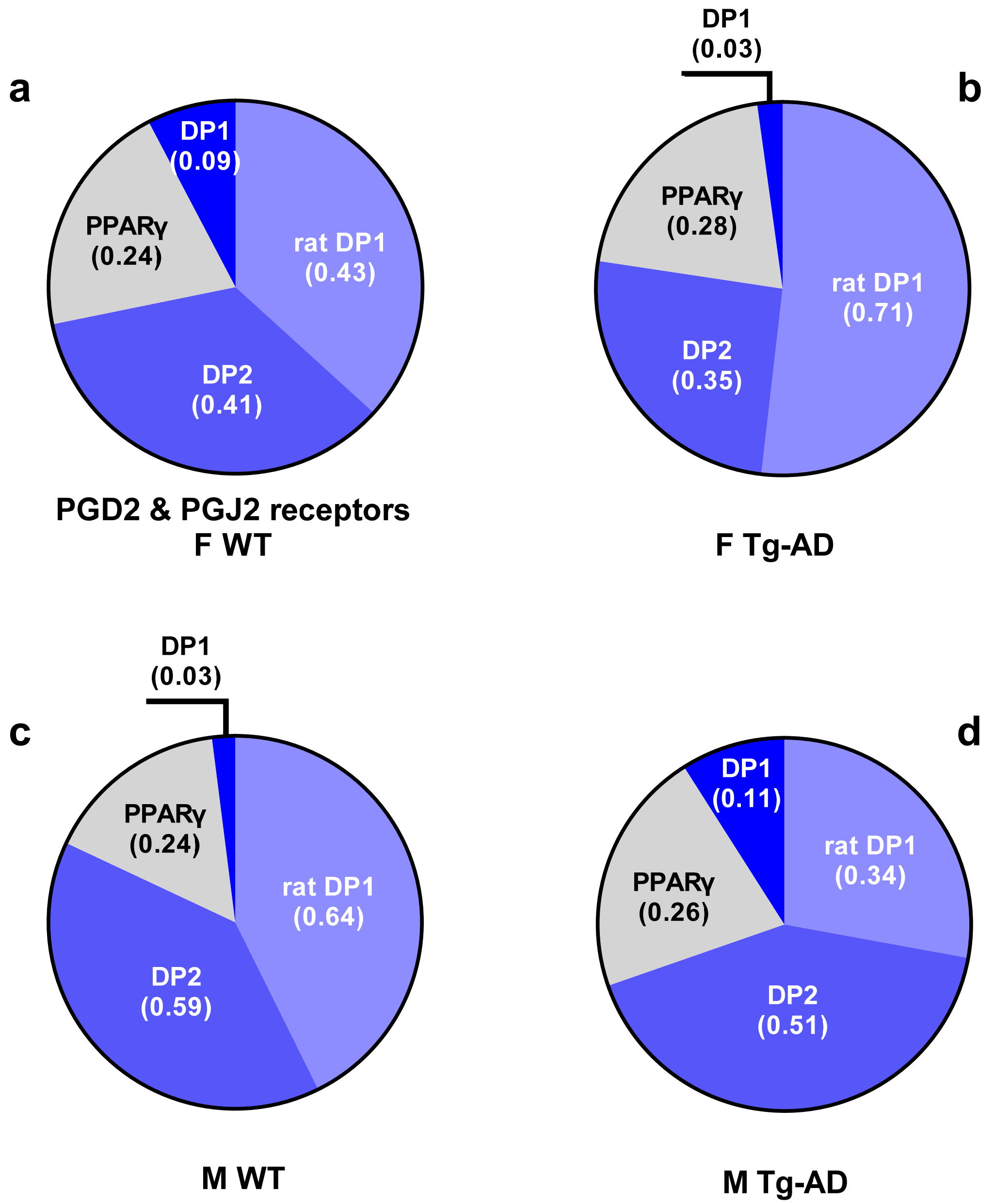

#### Supplemental Fig. 5

RNAseq PPARγ receptor activators & Sox-2

**a**

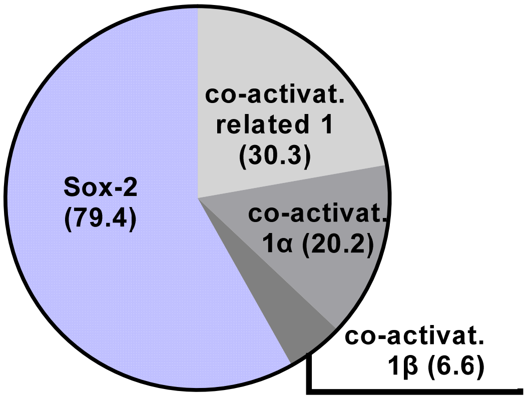


**PPARγ receptor activators**
**& PGD2 target**
**F WT**

**b**

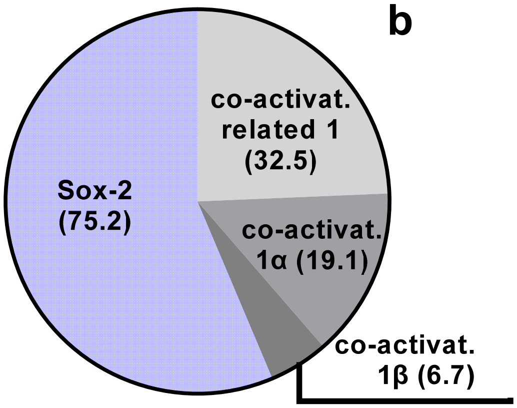


**F Tg-AD**

**c**

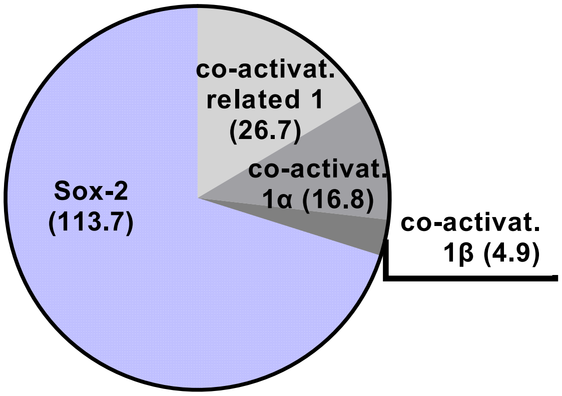


**M WT**

**d**

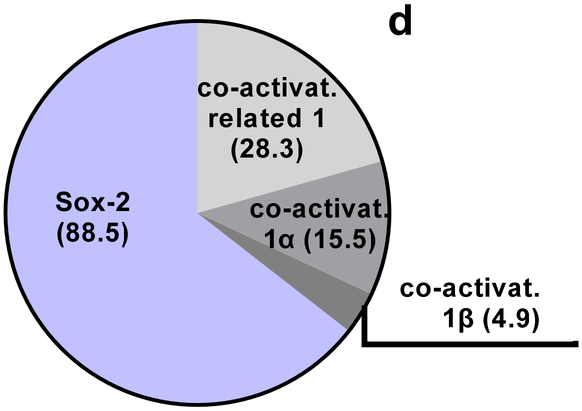


**M Tg-AD**

**Supplemental Fig. 6**

RNAseq – PGE2 receptors

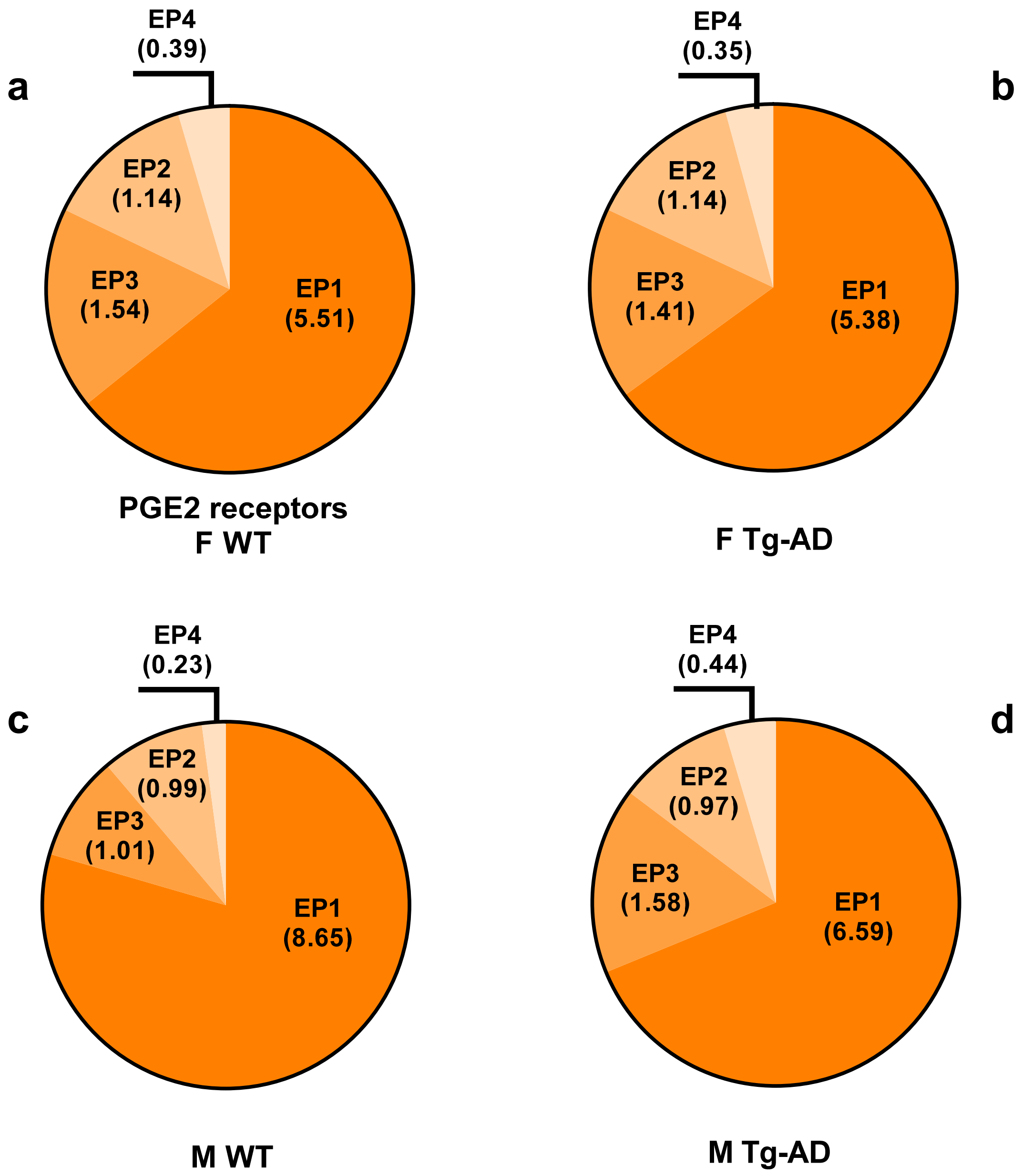

**Supplemental Fig. 7**
Western blot analyses

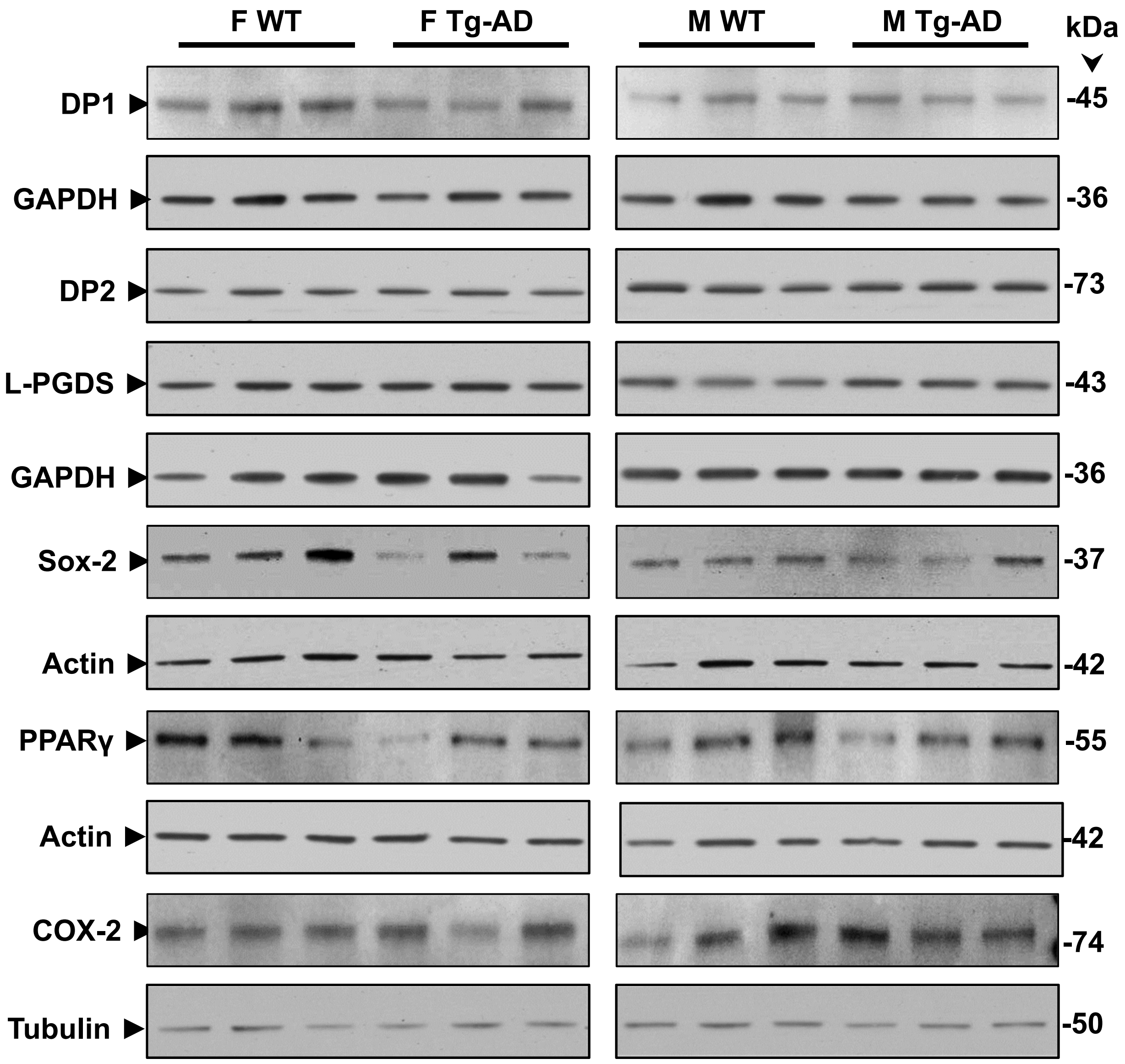

#### Supplemental Fig. 8
Full blots – DP1, DP2, L-PGDS

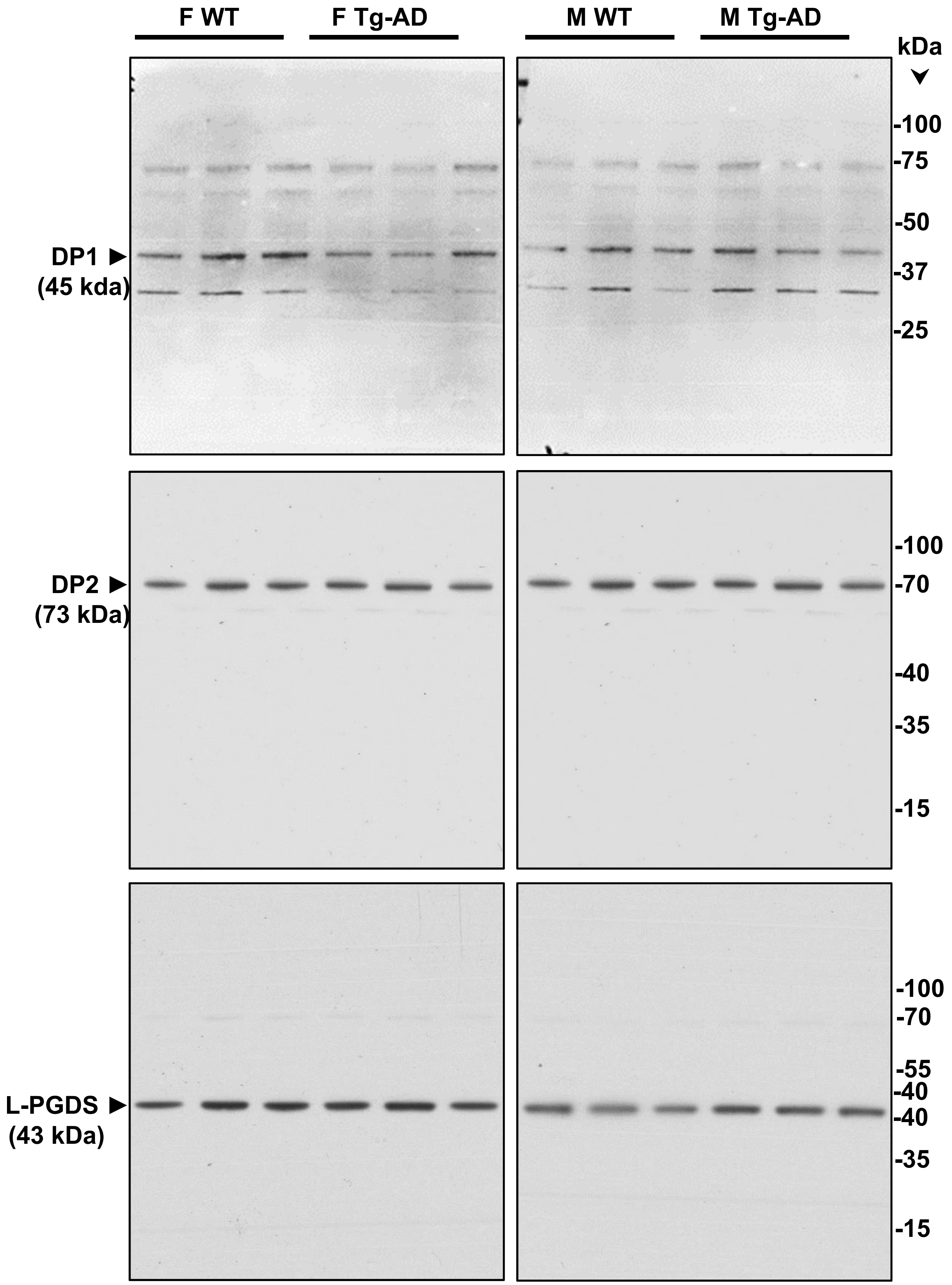

#### Supplemental Fig. 9

Full blots – FL-APP and Aβ

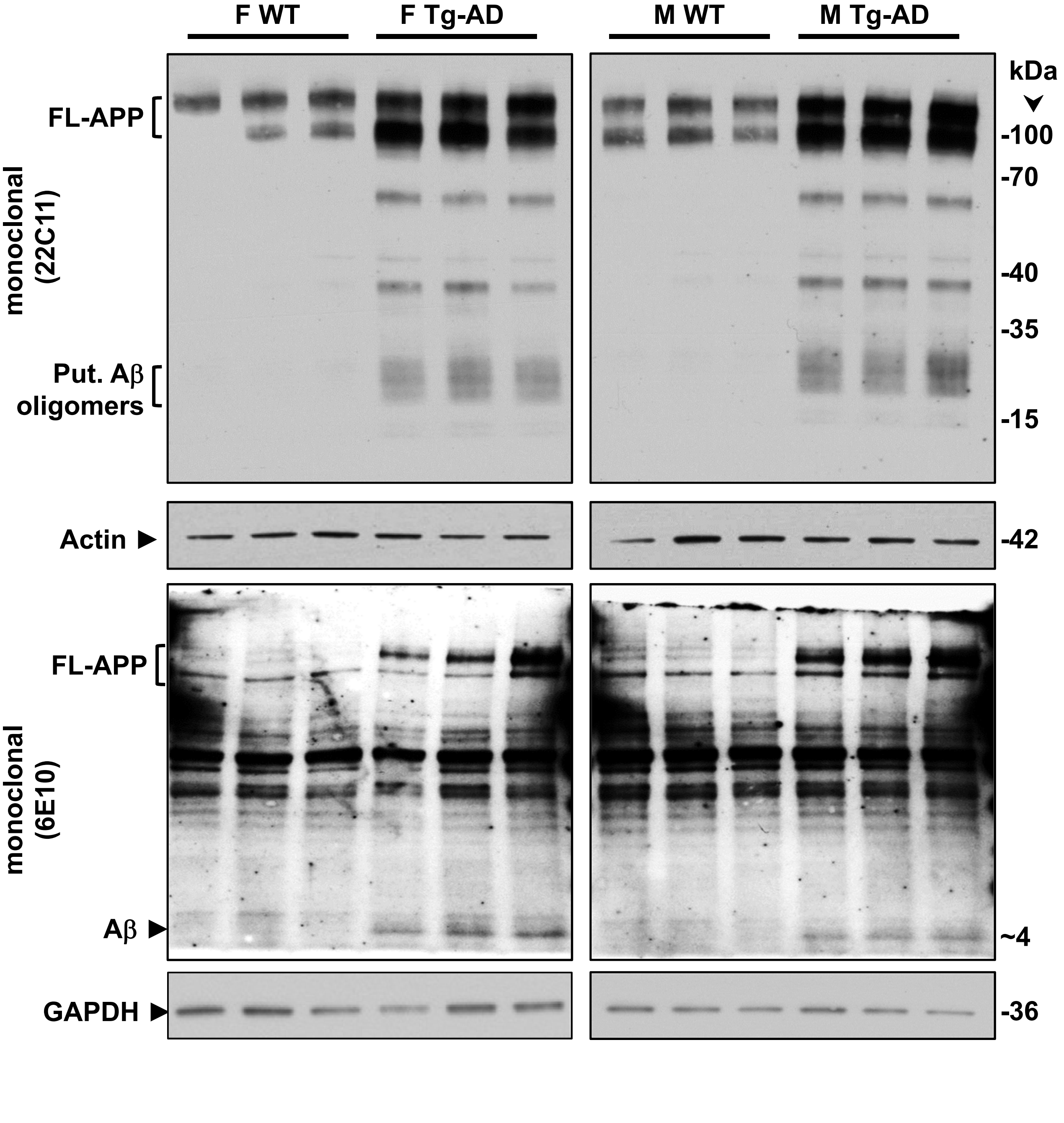

**Supplemental Fig. 10A**

IHC analysis of the right dorsal hippocampus of Tg-AD (left column, n = 9) and timapiprant-treated Tg-AD (right column, n = 9). 500 μm scale bars. DP1 (red), microglia (green, Iba1 antibody), and DP1/microglia co-localization (yellow).

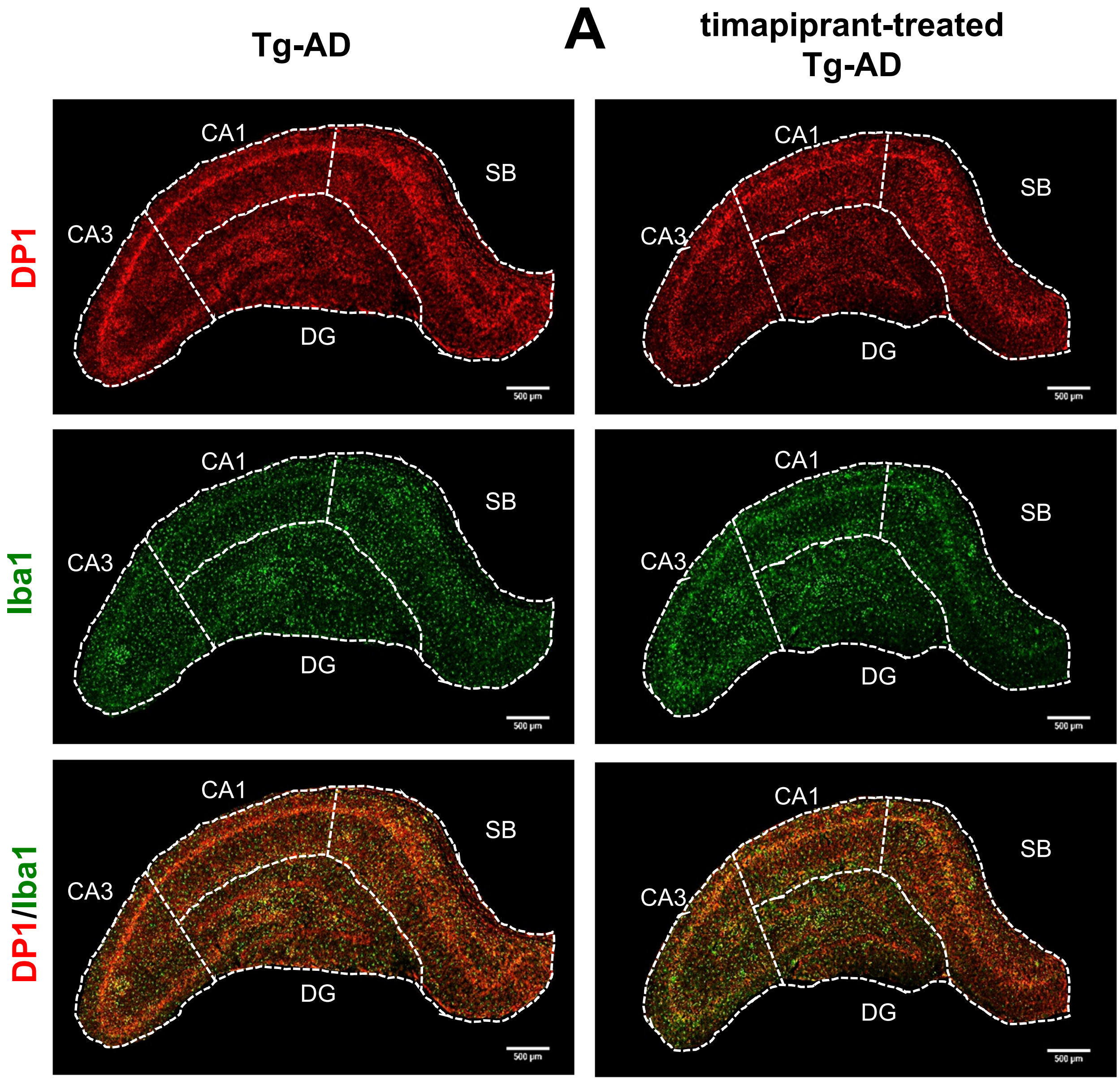

**Supplemental Fig. 10B**

IHC analysis of the right dorsal hippocampus of Tg-AD (left column, n = 9) and timapiprant-treated Tg-AD (right column, n = 9). 500 µm scale bars. DP2 (red), neurons (green, NeuN antibody), and DP2/neuronal co-localization (yellow).

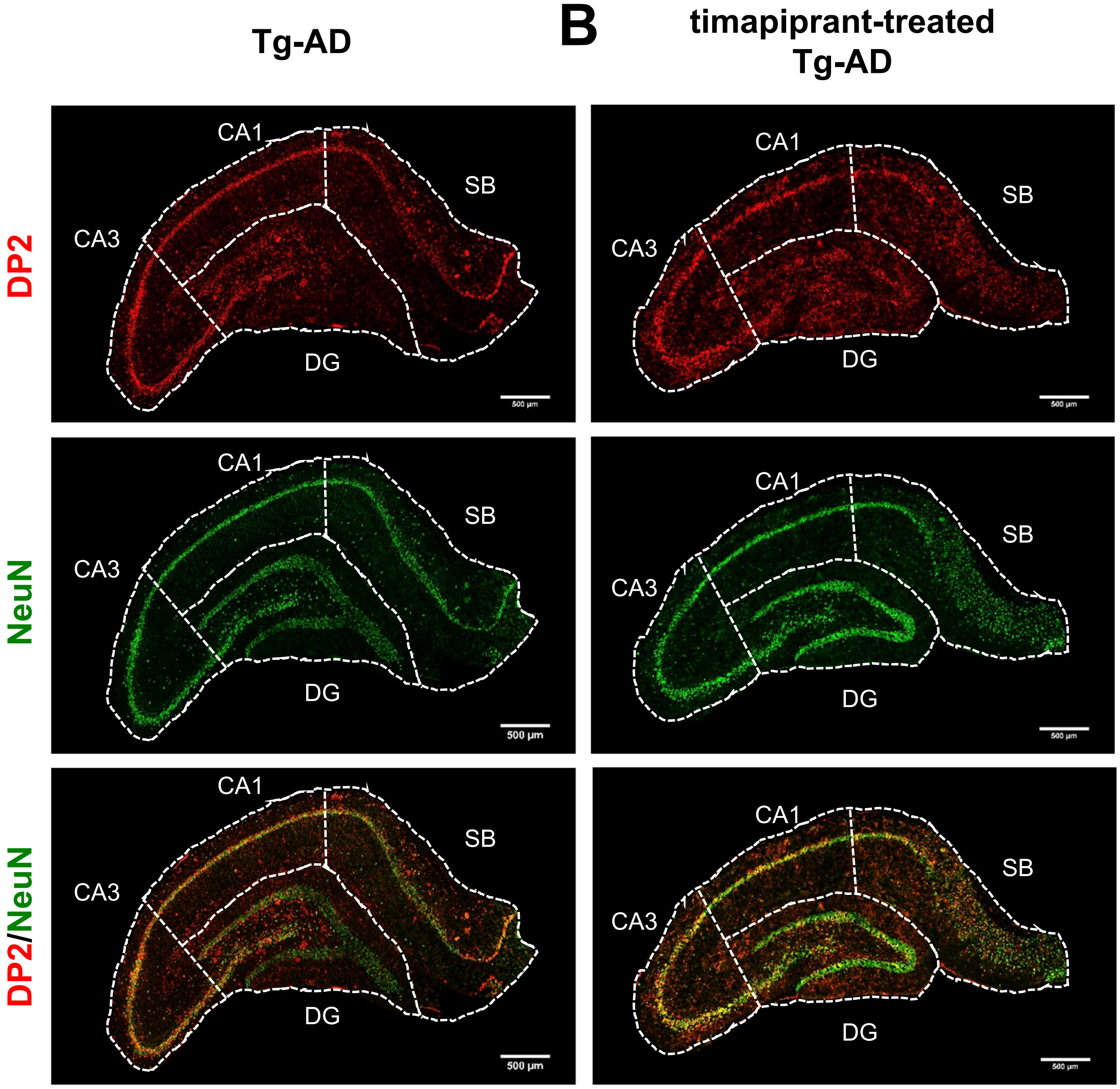

**Supplemental Fig. 10C**

IHC analysis of the right dorsal hippocampus of Tg-AD (left column, n = 9) and timapiprant-treated Tg-AD (right column, n = 9). 500 µm scale bars. Immunohistochemistry for Aβ (green) plaque load.

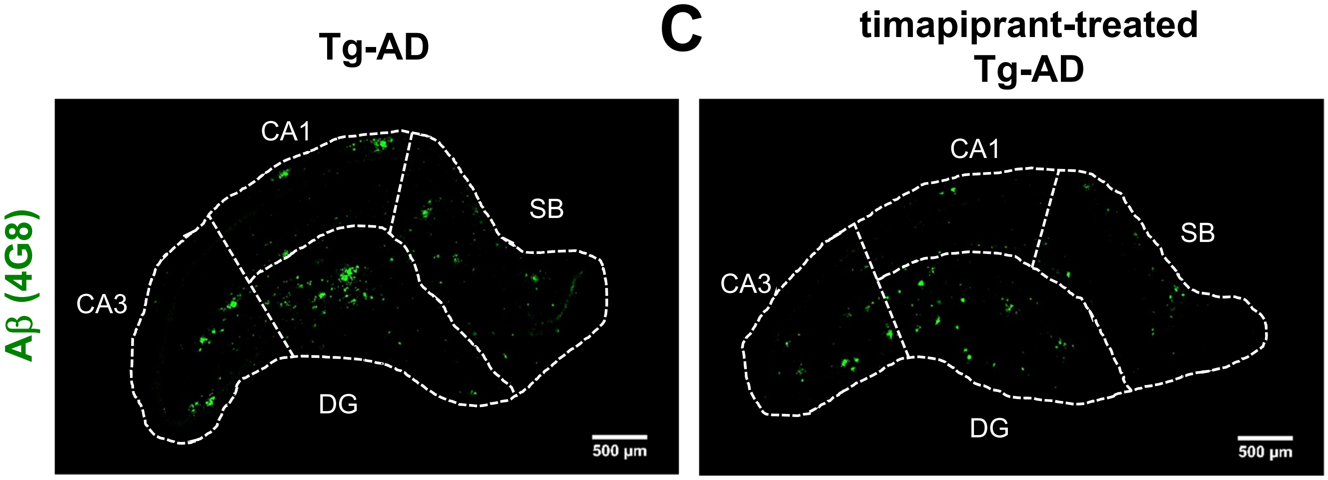
